# *Plasmodium falciparum* egress disrupts endothelial junctions and activates JAK-STAT signaling in a microvascular 3D blood-brain barrier model

**DOI:** 10.1101/2024.10.15.618439

**Authors:** Livia Piatti, Alina Batzilla, Fumio Nakaki, Hannah Fleckenstein, François Korbmacher, Rory K.M. Long, Daniel Schraivogel, John A. Hawkins, Tais Romero-Uruñuela, Borja López-Gutiérrez, Silvia Sanz, Yannick Schwab, Lars M. Steinmetz, James Sharpe, Maria Bernabeu

## Abstract

Cerebral malaria is a severe neurovascular complication of *Plasmodium falciparum* infection, with high mortality rates even after treatment with effective antimalarials. Limitations in current experimental models have hindered our knowledge of the disease. We developed a 3D blood-brain barrier (BBB) model with enhanced barrier properties using primary brain endothelial cells, astrocytes and pericytes. Exposure to parasite egress products increased microvascular permeability, likely due to transcriptional downregulation of junctional and vascular development genes in endothelial cells. In addition, it increased the expression of ferroptosis markers, antigen presentation and type I interferon genes and upregulated the JAK-STAT pathway across all BBB cell types. Incubation with cytoadherent schizont-stage *P. falciparum*-infected erythrocytes induced a similar, but highly localized transcriptional shift, along with inter-endothelial gaps at sites of parasite egress, significantly increasing permeability. The co-administration of egress products with the JAK-STAT inhibitor Ruxolitinib prevented junctional disruption and BBB breakdown. These findings provide key insights into the parasite-mediated mechanisms driving brain microvascular pathogenesis in cerebral malaria and suggest potential avenues for adjunctive therapies.

## Main

*Plasmodium falciparum* infections account for the majority of the 600,000 annual malaria deaths worldwide^1^, with cerebral malaria (CM) being one of the deadliest complications. Histological examination of post-mortem brain samples from CM patients has identified the sequestration of *P. falciparum-*infected red blood cells (iRBC) in the brain microvasculature as a key disease hallmark, often accompanied by vascular pathology and endothelial dysfunction^2–4^. Recent magnetic resonance imaging (MRI) studies suggest that fatal brain swelling in pediatric CM patients likely results from blood-brain barrier (BBB) dysfunction, disrupting the selective transport of fluids and molecules from blood vessels to the brain parenchyma and leading to vasogenic edema^5,6^. Even with treatments that rapidly clear parasites from blood^7^, CM still has a 15-20% mortality rate^8^, and half of the survivors suffer from long-term neurological and behavioral sequelae^9,10^. A better mechanistic understanding of how *P. falciparum* disrupts the BBB is crucial for developing adjunctive host-targeted treatments that could prevent deaths and long-term disabilities.

Two key parasite-induced disruptive mechanisms have been proposed. The first involves parasite binding to endothelial receptors endothelial protein C receptor (EPCR)^11^ and intercellular adhesion molecule 1 (ICAM-1)^12–14^, and blockade of EPCR homeostatic functions^15,16^. This pathogenic mechanism is exclusive to *P. falciparum* and does not occur in other malaria species, including rodent malaria models. The second mechanism suggests that parasites egressing from iRBC release endothelial-disruptive products, such as heme^17^, hemozoin^18^, parasite histones^19,20^, *P. falciparum* histidine-rich protein 2 (PfHRP2)^21^, or glycophosphatidyl inositol (GPI)^22^. Previous studies on these two mechanisms have been conducted using *in vitro* endothelial-only cultures with reduced barrier properties. Nevertheless, the BBB is a multicellular interface whose properties arise from the physical and cellular cross-talk between endothelial cells, pericytes and astrocytes.

Furthermore, mechanical cues from blood flow and the extracellular matrix enhance its barrier function^23^. Here, we have developed a bioengineered microvascular model that incorporates all these components, resulting in improved barrier properties. The use of this model has revealed that parasite products released during egress are responsible for an increase in vascular permeability, likely as a consequence of downregulation of endothelial junctional and vascular development pathways. We further demonstrated that *P. falciparum* egress products elicit the activation of inflammatory, including JAK-STAT, and antigen presentation pathways in all the cells that compose the BBB. When experiments were performed with cytoadhesive *P. falciparum-*iRBC, the disruptive effects were locally confined to regions of high sequestration and parasite egress. Yet, they still led to an increase in BBB permeability. Ruxolitinib, an inhibitor of the JAK-STAT pathway, prevented *P. falciparum-*induced barrier leakage, highlighting a link between inflammatory and vascular disruptive pathways.

## Results

### Primary brain astrocytes and pericytes improve endothelial barrier function

To determine the barrier disruptive pathways of *P. falciparum* in the brain microvasculature, we developed a bioengineered 3D-BBB model. The microvascular model was fabricated in a type I collagen hydrogel, pre-patterned by a combination of soft lithography and injection molding that generates a microfluidic 13 x 13 grid^24,25^. Commercial primary human astrocytes and brain vascular pericytes were seeded in the bulk collagen solution, and primary human brain microvascular endothelial cells were seeded under gravity-driven flow into the microfluidic network (Fig. 1a). The identity of the three cell types was confirmed by the expression of specific markers including platelet endothelial cell adhesion molecule 1 (PECAM-1), von Willebrand factor (vWF), vascular endothelial (VE)-cadherin and β-catenin for endothelial cells, platelet-derived growth factor receptor β (PDGFRβ) and nerve/glial antigen 2 (NG2) for pericytes and glial fibrillary acidic protein (GFAP), S100B and aquaporin 4 (AQP4) for astrocytes (Extended data Fig. 1a). After two days in culture, endothelial cells line the perfusable microvessels and are surrounded by a second layer of astrocytes and pericytes that sparsely contact the endothelium, resembling the 3D organization and architecture of the *in vivo* BBB (Fig. 1b). Generally, pericytes and astrocytes ensheath the abluminal side of microvessels, and collagen-residing astrocytes occasionally extend their end-feet towards endothelial cells (Fig. 1c,d). Quantitative RT-PCR measurements comparing two different astrocyte-to-pericyte ratios revealed that co-culture with astrocytes and pericytes at a 7:3 ratio increased the expression of endothelial BBB-specific markers over time, including tight junction markers (*OCLN, CLDN5*), BBB-transporters (*LRP1, SLC* family) and efflux pumps (*PGP, ABC* family), peaking after 7 days in culture (Extended data Fig. 1b). Notably, the increased expression of BBB markers was associated with an improvement in microvascular barrier properties. The 3D-BBB model containing pericytes and astrocytes showed a significant decrease in permeability to 70 kDa FITC-dextran (2.15x10^-6^ cm/s – Interquartile range (IQR) = 0.30, 6.37), compared to an endothelial-only model (8.31x10^-6^ cm/s – IQR = 3.98, 12.40) (Fig. 1e and Extended data Fig. 1c). Thus, the enhanced permeability properties of our bioengineered 3D-BBB model provide a unique opportunity to study the mechanisms of vascular barrier disruption induced by *P. falciparum*.

**Figure 1.**
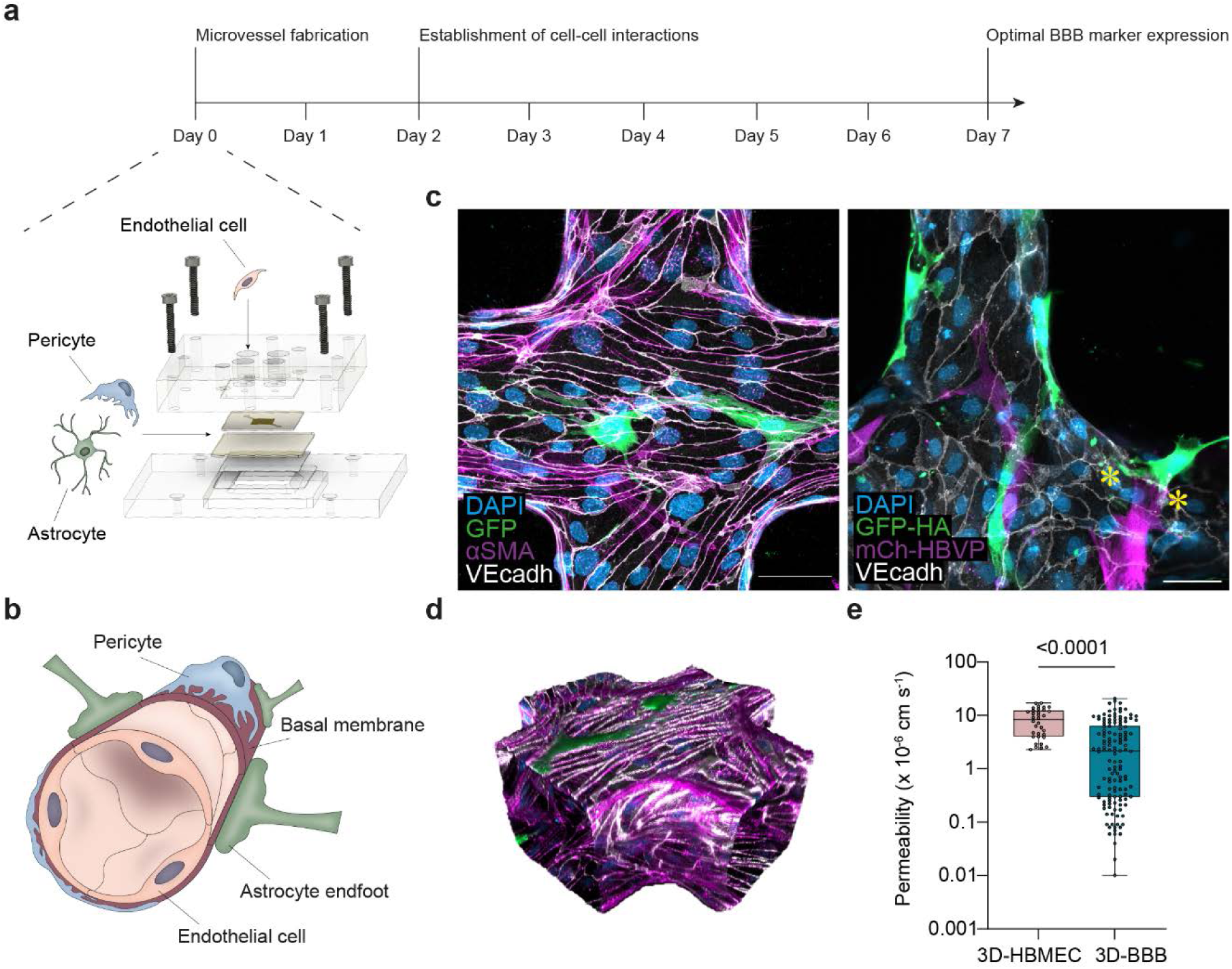
Perfusable 3D-BBB microvascular model recapitulates the BBB structure, cell identities and presents improved barrier properties. **a**, Representation of the experimental timeline for 3D-BBB microvessel fabrication and culture, including a schematic sketch showing the materials for microvessel fabrication and the cells that compose it: human primary astrocytes, pericytes and brain microvascular endothelial cells. b, Schematic depiction of the BBB architecture. c, Representative maximum z-projection images of the 3D-BBB microvessel model and the spatial multicellular organization of endothelial cells stained with VE-cadherin (white), GFP-expressing astrocytes and pericytes labeled with αSMA (pink) (left image) or expressing mCherry (right image). Asterisks indicate astrocyte end feet contacting endothelial cells. d, 3D reconstruction of a portion of the microfluidic network. e, Apparent microvascular permeability to 70 kDa FITC-dextran. Each point represents an ROI from endothelial-only (N = 3) and 3D-BBB (N = 13) microvessels (Mann-Whitney U test).

### *P. falciparum*-iRBC products released during egress downregulate endothelial junction expression and increase microvascular permeability

The egress of *P. falciparum-*iRBC has long been recognized as an endothelial disruptive event^18–22,26^. However, previous studies were done in models with a weaker vascular barrier and did not include astrocytes or pericytes^27–30^. We first explored how parasite egress products contribute to malaria pathogenesis within our newly engineered 3D-BBB model. To this end, we generated a solution containing *P. falciparum* products released upon the egress of 5x10^7^ tightly synchronized iRBC/mL (hereafter referred to as iRBC-egress media)^31^ (Extended Data Fig. 2a). We estimate that this concentration of products is equivalent to 5x10^4^ parasites/μL, levels often found in CM patients^5^. 3D-BBB microvessels were incubated with iRBC-egress media for 24 hours and subjected to a multimodal analysis. This included single-cell RNA sequencing (scRNA-seq) on dissociated microvessels, electron and confocal microscopy on fixed microvessels, as well as live permeability measurements (Fig 2a). As a control, we perfused the supernatant of an uninfected erythrocyte (uRBC) control processed in the same way as the infected counterpart.

**Figure 2.**
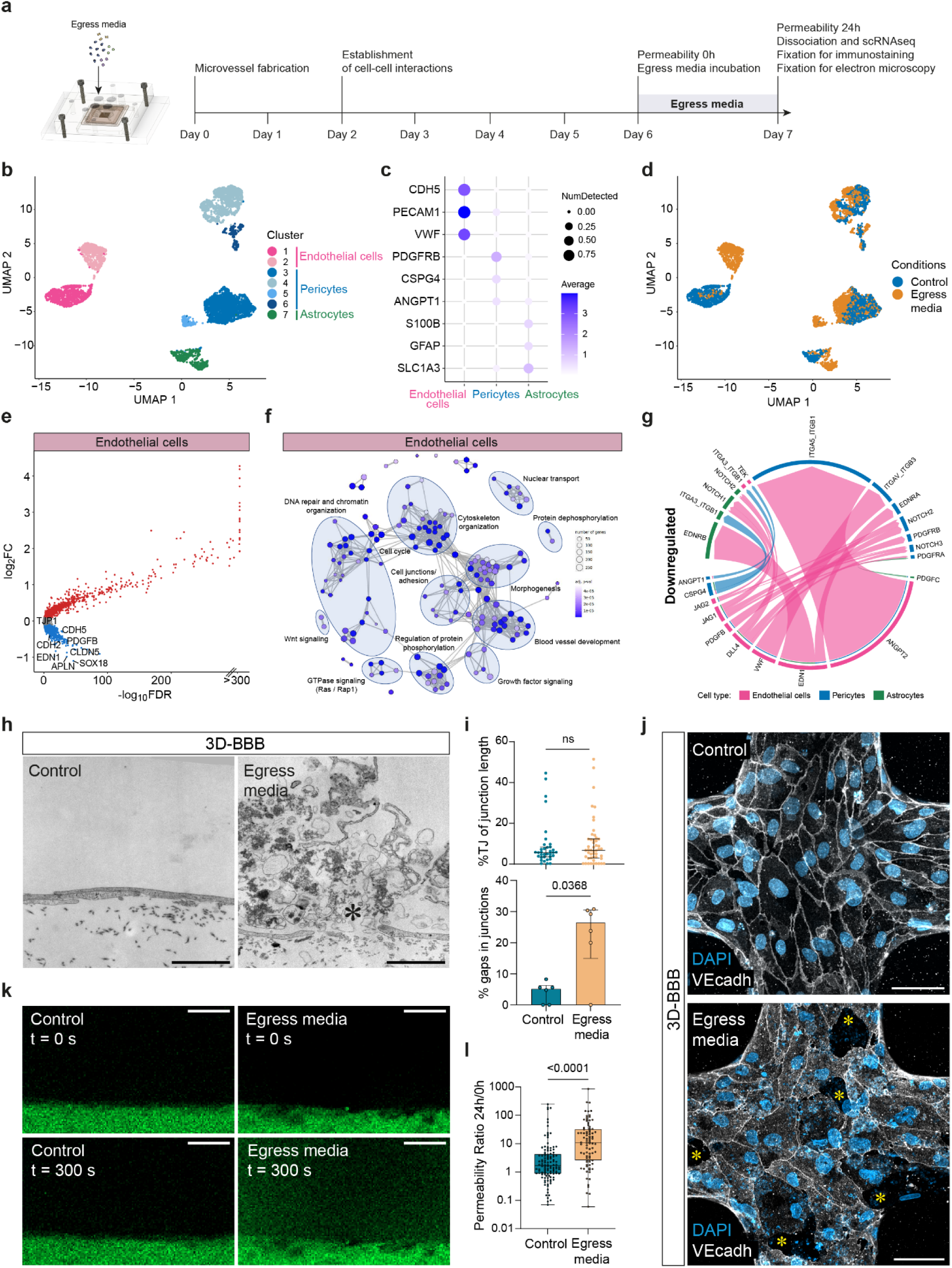
iRBC-egress media induces transcriptional downregulation of junctional markers and BBB signaling, and causes inter-endothelial gaps impairing microvascular integrity. **a,** Representation of the experimental timeline on 3D-BBB microvessels before and after 24-hour incubation with iRBC-egress media **b-g**, Single-cell transcriptomic analysis comparing 3D-BBB models exposed to iRBC-egress media and control uRBC media. **b**, UMAP of sequenced cells colored by unsupervised *Leiden* clustering. **c,** Dot plot of main BBB cell type markers. **d,** UMAP of sequenced cells colored by experimental condition. **e**, Volcano plot of differentially expressed genes in endothelial cells upon 24-hour iRBC-egress media incubation, plotting the log2-transformed fold change (log2FC) against the statistical significance (-log10 of the false discover rate (FDR)). Significantly up- or downregulated genes (FDR<0.05, log2FC>0.1 or log2FC<-0.1, respectively) are marked in red or blue and selected downregulated genes are labeled. **f**, GO-term over-representation analysis on significantly downregulated genes (FDR<0.05, log2FC< -0.1) in endothelial cells. Each network node represents one of the most significant GO-terms (adjusted p-value < 0.0001) and edges connect GO-terms with more than 20% gene overlap. GO-term clusters were manually summarized with one label term. **g**, Selected downregulated ligand-receptor interactions important for BBB-establishment, identified among the three BBB cell types after exposure to iRBC-egress media using the *CellChat* package. Arrows point from ligands on sender cells to receptors on receiver cells and are colored by sender cell. Weights of links are proportional to the interaction strength. **h,** TEM images showing an inter-endothelial gap formed in 3D-BBB microvessels exposed to iRBC-egress media (right), compared to an intact junction in control microvessels (left). Asterisk indicates parasite egress products. Scale bar = 2 μm. **i**, Scatter plot comparing the percentage of tight junctions over total junctional length in 3D-BBB microvessels exposed to control uRBC media or iRBC-egress media (Mann-Whitney U test) (top), and bar plot quantifying the percentage of inter-endothelial gaps found by TEM on 3D-BBB microvessels incubated with control uRBC media (N = 1) or iRBC-egress media (N = 2) (bottom). **j**, Maximum z-projection of a confocal image showing junctional marker VE-cadherin (white) in a microvessel incubated with control media or iRBC-egress media. DAPI (blue) stains large nuclei corresponding to BBB cells, and nucleic acids found in iRBC-egress media, presenting small punctate labeling. Asterisks indicate inter-endothelial gaps. Scale bar = 50 μm. **k**, Representative images showing 70 kDa FITC-dextran diffusion through the lateral wall of the 3D-BBB before and after 24-hour incubation with control media or iRBC-egress media. Scale bar = 50 μm. **l**, Ratio of the apparent permeability calculated at 24-hour post-incubation and at pre-incubation with control media or iRBC-egress media. Each point represents the ratio from an ROI coming from 3D-BBB microvessels exposed to control media (N = 4) and iRBC-egress media (N = 10) (Mann-Whitney U test).

The 3D-BBB microvessels were disassembled and dissociated into a single-cell suspension, followed by sample-multiplexing using the MULTI-seq protocol^32^ to pool infected and control conditions for scRNA-seq. The 6454 quality-controlled cells in the resulting scRNA-seq dataset were visualized using a uniform manifold approximation and projection (UMAP) algorithm, revealing 7 distinct clusters after unsupervised clustering (Fig. 2b). Cell type annotation showed the presence of two clusters with high expression of endothelial markers (cluster 1 and 2: *CDH5, VWF* and *PECAM1*), 4 clusters expressing pericyte markers (clusters 4 and 6: high expression of *PDGFRB,* and clusters 3 and 5: moderate *PDGFRB*) and a cluster expressing astrocytic markers (cluster 7: *S100B* and *GFAP)* (Fig. 2b,c and Extended Data Fig. 2b). A shift in the transcriptional profile of the three cell types was observed upon exposure to iRBC-egress media, with endothelial cells splitting into a separate cluster from the control population (Fig. 2d). Differential gene expression analysis on endothelial cells revealed that iRBC-egress media caused a significant decrease in the expression of multiple genes encoding proteins that are critical for BBB integrity, including the tight junction genes *CLDN5* (Claudin-5) and *TJP1* (ZO-1) and the adherens junction transcript *CDH5* (VE-cadherin) (Fig. 2e and Supplementary Information Table 1). Gene ontology term (GO-term) over-representation analysis indicated that most downregulated transcripts in endothelial cells are associated with endothelial cell junctions and adhesion, cytoskeleton organization, blood vessel development, DNA repair and chromatin organization and Wnt signaling (Fig. 2f and Supplementary Information Table 2). We employed the *CellChat* package^33^ to analyze differential ligand-receptor interactions among the three cell types present in the model after exposure to iRBC-egress media. The analysis showed a decrease in vascular signaling interactions (*VWF, EDN1*), as well as in important pathways for endothelial-pericyte homeostasis, including signaling of angiopoietin-1 (*ANGPT1*), PDGF-BB (*PDGFB),* and CSPG4 (*NG2*). Additionally, iRBC-egress media caused a decrease in key endothelial-pericyte-astrocyte signaling molecules, including the Notch pathway (*DLL4, JAG1/2*) (Fig 2g and Extended Data Fig. 3). Overall, exposure to iRBC-egress media led to the downregulation of transcripts associated with endothelial integrity and impaired major signaling pathways among all cell types present in the model.

To test whether these transcriptional changes resulted in an impairment of 3D microvascular integrity, we first quantified the number of inter-endothelial gaps, as well as the percentage of junctional length covered by electron-dense tight junctions by transmission electron microscopy (TEM). Parasite egress products were visible by TEM and appeared as fuzzy aggregates containing parasite organelles and hemozoin, as well as parasitophorous and iRBC membranes with knobs (Fig. 2h). Although we observed a significantly higher percentage of ultrastructural gaps in 3D-BBB microvessels exposed to iRBC-egress media, we did not observe any significant differences in the length of tight junctions in regions of cell-cell contact that remained intact (Fig. 2i). Changes in junction morphology were also observed by confocal microscopy. Specifically, VE-cadherin labelling revealed an altered adherens junction pattern compared to the control condition, with thin junctional staining and the formation of large inter-endothelial gaps (Fig. 2j). The transcriptional and morphological changes observed were accompanied by functional changes in vascular barrier. Baseline microvascular permeability was measured by 70 kDa FITC-dextran perfusion on day 6 of 3D-BBB microvessel formation, followed by a second measurement in the same regions of interest 24 hours after addition of iRBC-egress or control media (Fig. 2k). After 24 hours, 3D-BBB microvessels treated with iRBC-egress media presented a significant 6.5-fold increase in microvascular permeability ratio (11.07 – IQR = 2.60, 32.56) compared to controls (1.68 – IQR = 0.87, 4.28) (Fig. 2l). Notably, no significant differences in microvascular permeability ratio were observed between the two HBMEC donors used in this study (Extended data Fig. 2c), nor in 3D-BBB microvessels incubated with uRBC media control (Extended Data Fig. 2d). These results suggest that *P. falciparum-*iRBC products decrease the expression of tight and adherens junction markers, changing the morphology of inter-endothelial junctions, together with a functional impairment of vascular barrier integrity.

### *P. falciparum* egress products induce a global activation of inflammatory, JAK-STAT, antigen presentation and ferroptosis-associated pathways in the 3D-BBB model

Studies in *P. berghei* rodent CM models have shown the ability of endothelial cells to cross-present parasite antigens present in merozoites^34,35^. Our analysis revealed that *P. falciparum* could induce a similar behavior in all the cells that compose our bioengineered BBB model. Specifically, exposure of 3D-BBB microvessels to iRBC-egress media caused the upregulation of multiple genes associated with inflammatory and antigen presentation pathways in endothelial cells (Fig. 3a). The same upregulated transcripts were identified in pericytes and astrocytes, suggesting that iRBC-egress media has the potential to cross the endothelial barrier. We found an upregulation of transcripts involved in type I interferon (IFN) response and anti-viral pathways, including genes of the IFN-induced protein with tetratricopeptide repeats (IFIT) gene family (e.g. *IFIT1*, *IFIT2*, *IFIT3)*, IFN-stimulated genes (ISGs) (e.g. *ISG15, ISG20*), and other IFN-inducible genes (e.g. *MX1*, *IFI6, IFI27, OAS1),* as well as ferroptosis genes (e.g. *HMOX1*) (Fig. 3a). Furthermore, transcripts upregulated by *P. falciparum* egress products include members of the JAK-STAT family of signal transducers (*STAT1*, *STAT2*, *JAK2*), a signaling pathway that induces the expression of ISGs, and some of their interacting proteins (*IRF1, IRF9*). GO-term over-representation analysis on the significantly upregulated transcripts confirmed a global increase in expression of transcripts associated with cytokine-, viral-, and type I IFN response, as well as antigen presentation, NFκB signaling, and protein catabolism in all cell types that compose the BBB (Fig. 3b and Supplementary Information Table 2). Cell type-specific processes that were significantly upregulated include apoptosis and autophagy in endothelial cells and astrocytes, ER stress and Golgi/vesicle transport in endothelial cells and regulation of cell migration in pericytes (Fig. 3b and Supplementary Information Table 2). *CellChat* analysis on upregulated transcripts revealed an increase in the expression of inflammatory ligand-receptor pairs following exposure to iRBC-egress media. Pericytes and astrocytes showed an increased expression of collagen encoding genes, suggestive of a fibrotic, scar-forming phenotype (Fig. 3c and Extended Data Fig 3). Additionally, all three cell types showed elevated expression of midkine (*MDK*), a chemoattractant for the recruitment of neutrophils, macrophages and lymphocytes^36^.

**Figure 3.**
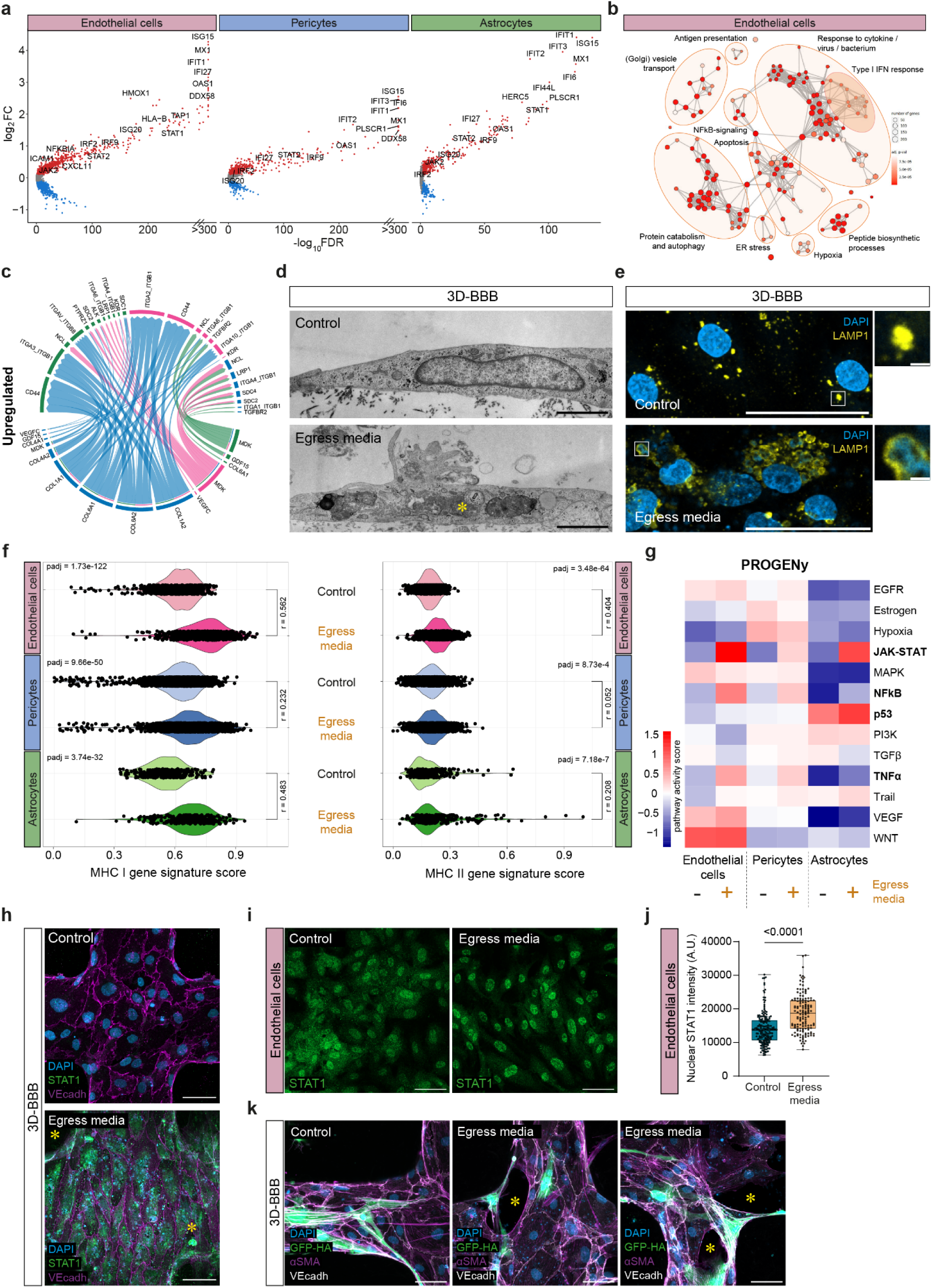
iRBC-egress media activates inflammatory and antigen presentation pathways in all BBB cell types. **a**, Volcano plots of differentially expressed genes in endothelial cells (same analysis as in 3e), pericytes, and astrocytes upon 24-hour incubation with iRBC-egress media plotting the log2-FC against the statistical significance (-log10 of the FDR). Significantly up- or downregulated genes (FDR<0.05, log2FC>0.1 or log2FC<-0.1, respectively) are marked in red or blue and selected upregulated genes are labeled. **b**, GO-term over-representation analysis on significantly upregulated genes (FDR<0.05, log2FC>0.1) in endothelial cells. Each network node represents one of the most significant GO-terms (adjusted p-value < 0.0001) and edges connect GO-terms with more than 20% gene overlap. GO-term clusters were manually summarized in one label term. **c**, Selected upregulated ligand-receptor interactions identified among the cell types within the 3D-BBB model after exposure to iRBC-egress using the *CellChat* package. Arrows point from ligands on sender cells to receptors on receiver cells and are colored by sender cell. Weights of links are proportional to the interaction strength. **d**, Representative TEM images of endothelial cells within the 3D-BBB model taking up parasite material (asterisk) after incubation with iRBC-egress media (bottom) and representative control endothelial cell incubated with uRBC media (top). Scale bar = 2 μm. **e**, Representative confocal imaging (left) and close-up views (right) showing maximum z-projection of LAMP1 labeling (yellow) in 3D-BBB microvessels after 24-hour incubation with iRBC-egress media or control media. Nuclei and *P. falciparum* DNA were labelled with DAPI (blue). Scale bar = 50 μm; close-up scale bar = 2 μm. **f**, Violin plots showing the MHC I and MHC II gene signature score in every cell plotted by cell type and condition (Mann-Whitney U test followed by calculation of effect size r). Genes in the MHC I and II signature score can be found in the Methods. **g**, Heatmap of pathway activities inferred using the *PROGENy* method. The colors in the heatmap correspond to the mean PROGENy pathway activity score in the cells of the respective cell type and condition. **h**, Representative confocal imaging showing maximum z-projection of STAT1 labeling (green) in 3D-BBB microvessels after 24-hour incubation with iRBC-egress media or control media. Asterisks indicate STAT1-positive astrocytes and pericytes in the collagen hydrogel. Endothelial junctions were labeled with VE-cadherin (magenta) and nuclei with DAPI (blue). Scale bar = 50 μm. **i**, Mean fluorescence intensity of STAT1 labeling in the nuclei of endothelial cells grown on monolayers (N = 6/condition) after 24-hour incubation with iRBC-egress media or media control (Mann-Whitney U test). **j**, Representative maximum z-projection of a confocal image showing STAT1 protein localization (green) in endothelial monolayers from the quantification in j,. Scale bar = 50 μm. **k**, Representative confocal imaging showing maximum z-projection of 3D-BBB microvessels after 24-hour incubation with iRBC-egress media. GFP-expressing astrocytes (green) and αSMA-labelled pericytes (magenta) asterisks represent gaps between endothelial cells, stained with VE-cadherin (white). Scalebar = 50 μm.

Antigen presentation appeared to be an important inflammation-related process that was strongly elevated in all cell types that compose the 3D-BBB model. Notably, we found evidence of cellular uptake of parasite material by TEM. Endothelial cells in the 3D-BBB microvessels incubated with iRBC-egress media showed signs of activation, including the formation of membrane protrusions, large vacuoles containing iRBC membranes or hemozoin (Fig. 3d and Extended Data Figure 4a), indicating the activation of parasite uptake. Immunofluorescent staining shows colocalization of parasite nucleic acids with lysosome-associated membrane protein 1 (LAMP1) (Fig. 3e), suggesting the delivery of parasite material to the lysosomal compartment of endothelial cells. We therefore defined transcriptomic gene signatures for antigen presentation (see Methods), either for major histocompatibility complex (MHC) class I transcripts or MHC class II transcripts (Fig. 3f). Endothelial cells and astrocytes exhibited a robust upregulation of the MHC class I gene signature (effect size r > 0.5, p<0.0001), while only a modest upregulation was observed in pericytes (effect size r = 0.2, p<0.0001). Interestingly, the MHC class II antigen presentation gene signature, associated with CD4 T-cell recruitment, was also strongly upregulated in endothelial cells (effect size r = 0.4, p<0.0001) and modestly upregulated in astrocytes (effect size r=0.2, p<0.0001), with some astrocytes presenting particularly high MHC class II signature scores.

To gain deeper insights into other signaling pathways dysregulated upon exposure to *P. falciparum* egress products, we utilized PROGENy (Pathway RespOnsive GENes for activity inference)^37^, a computational method that infers signaling pathway activities based on downstream gene expression. Consistent with the GO-term analysis, we observed an increase in NFκB and TNFα signaling in endothelial cells and pericytes upon challenge with iRBC-egress media. Notably, the JAK-STAT pathway was activated across all cell types (Fig. 3g), a result consistent with the upregulation of transcripts associated with the type I IFN response (Fig. 3a, b). To validate this result, we performed immunofluorescent staining of STAT1 on cell monolayers and 3D-BBB microvessels. Increased STAT1 expression was observed in the 3D-BBB model upon 24-hour exposure to parasite egress products compared to the control condition, where the signal was barely detectable. In accordance with our scRNA-seq results, this increase occurred not only in endothelial cells, but also in the supporting pericytes and astrocytes present in the collagen hydrogel (Fig. 3h), indicating an overall response of the model to iRBC-egress media. STAT1 translocation to the nucleus was evaluated in 2D monolayers, as a proxy for increased pathway activity^38^. While endothelial cells presented increased STAT1 nuclear localization compared to the media-only condition, pericytes and astrocytes did not (Fig. 3i,j and Extended Data Fig. 4b,c). Furthermore, we found that the increase in the apoptosis-associated p53 pathway in astrocytes, as identified in the scRNA-seq analysis (Fig. 3g), was associated with a substantial reduction in cell density by immunofluorescence in astrocyte 2D monolayers (Extended Data Fig. 4d,e). This decrease occurred without an increase of astrocyte activation markers GFAP and ICAM-1 (Extended Data Fig. 4d). As a positive control for astrocytic activation, we treated astrocyte monolayers with TNFα, IL-1β and IFNγ or a cytokine cocktail at concentrations similar to those observed in CM patients^39^, which resulted in an increase in both GFAP and ICAM-1 expression in the absence of a significant reduction in cell density (Extended Data Fig. 4d,e). Despite the lack of astrocyte reactivity after exposure to *P. falciparum-*egress media, astrocytes were often found to extend their processes towards regions of endothelial disruption in our 3D-BBB model (Fig. 3k). Taken together, our results suggest that *P. falciparum* products released upon egress cause a significant upregulation of inflammatory and antigen presenting pathways, albeit with cell-specific differences.

### Binding of *P. falciparum*-iRBC for 6 hours induces minor transcriptional changes

Next, we aimed to investigate whether blockade of endothelial receptors such as EPCR and ICAM-1 by iRBC binding directly contributes to BBB pathogenesis, given its strong association with CM^6,16,40^. We perfused 3D-BBB microvessels with highly synchronized trophozoite (26-34 hours post infection) or schizont (38-46 hours post infection) stages of *P. falciparum* HB3var03, a parasite line expressing a dual EPCR-ICAM-1 binding PfEMP1. Parasites or uninfected RBC were perfused at the same concentration as iRBC*-*egress media (5x10^7^ iRBC/mL) for 30 minutes, followed by a 20-minute wash to release unbound iRBC. Microvessels were incubated for 6 hours, a timepoint that would prevent egress of parasites in the trophozoite condition, and analyzed morphologically through electron and confocal microscopy, as well as at the single-cell transcriptomic level (Fig. 4a). The UMAP confirmed the correct synchronization and development of *P. falciparum*-iRBC stages, as visualized by a trophozoite cluster positive for the *P. falciparum* mid-stage transcript *PfHB3_100020300* and a continuous, arch-shaped schizont cluster positive for late-stage marker *PfHB3_090035000* (Fig 4b and Extended data Fig 5a). The scRNA-seq dataset was then filtered to exclude *P. falciparum*-iRBC from the analysis to focus on the transcriptional changes in the BBB cell types. We obtained 4514 quality-controlled cells including all three conditions from the trophozoite, schizont and uninfected RBC microvessels. UMAP visualization after re-clustering of the BBB cells showed 4 distinct clusters, including an endothelial cluster (*CDH5* and *PECAM1*), an astrocyte cluster (*GFAP* and *S100B*) and two pericyte clusters (*PDGFRB^high^*and *PDGFRB^moderate^*) (Extended data Fig. 5a,b,c). In contrast to exposure to *P. falciparum*-iRBC egress media, the UMAP representation revealed no clear segregation of cells based on the experimental conditions (Fig. 4c). Nevertheless, we identified some dysregulated transcripts in endothelial cells that were similar to those observed in cells treated with iRBC-egress media (Fig. 4d). Incubation with both trophozoite and schizont stages led to the downregulation of endothelial tight junction marker *CLDN5*, as well as of its regulator *SOX18*. Exposure to trophozoite stages caused an upregulation of genes encoding for vesicle transport processes, including ER transcripts, such as *KDELR3*, or vesicular components, like *CAV1, COPB1* and *COPE* (Fig. 4d,e). Upon exposure to schizonts, we observed a downregulation of processes related to blood vessel development, including angiogenic, barrier formation- and endothelial migration-associated genes *APLN*, *END1, ENG, ROBO4* and *PDGFB* (Fig. 4e). Exposure to trophozoites and schizonts did not cause strong transcriptional changes in pericytes and astrocytes in the 3D BBB-model (Fig. 4f). Altogether, *P. falciparum-*iRBC cause minimal global transcriptional changes in human cells present in the 3D-BBB model.

**Figure 4.**
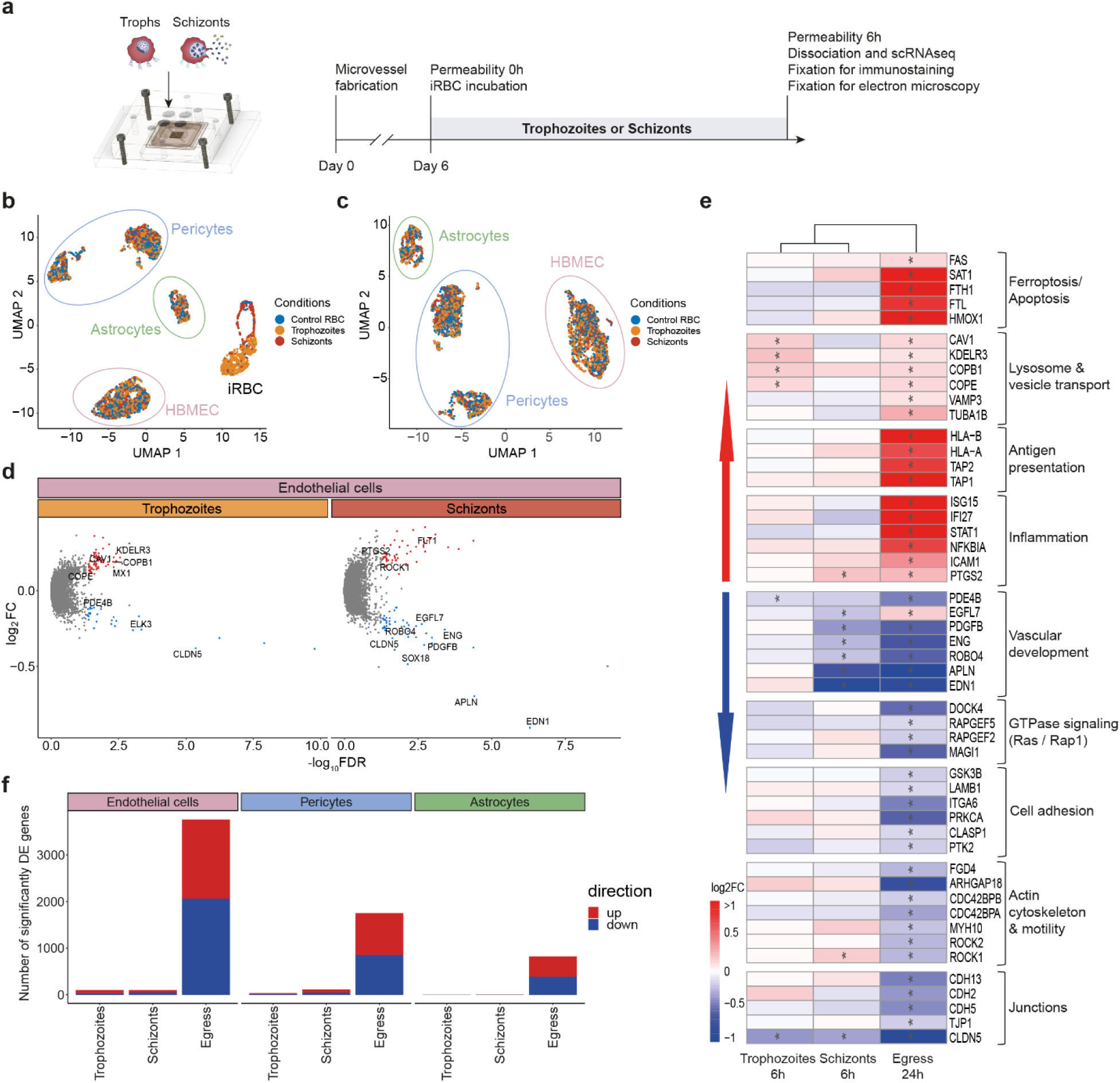
6-hour incubation with trophozoite- and schizont-stage *P. falciparum*-iRBC induces only minor transcriptional changes in BBB cell types. **a,** Representation of the experimental timeline on 3D-BBB microvessels before and after 6-hour incubation with cytoadherent trophozoite or schizont *P. falciparum*-iRBC or control devices perfused with uninfected RBC. **b**, UMAP of sequenced BBB cells and *P. falciparum*-iRBC colored by experimental condition. **c**, UMAP of BBB cells after exclusion of *P. falciparum*-iRBC. **d**, Volcano plots of differentially expressed genes in endothelial cells upon 6-hour incubation with trophozoite- or schizont-stage iRBC plotting the log2FC against the statistical significance (-log10 of the FDR). Significantly up- or downregulated genes (FDR<0.05, log2FC>0.1 or log2FC<-0.1, respectively) are marked in red or blue respectively, and selected genes are labeled. **e**, Heatmap showing log2FC values of selected genes belonging to significant GO-terms shown in Fig. 2f and 3b, in brain endothelial cells after 24-hour exposure to iRBC-egress media, or 6-hour exposure to trophozoite- or schizont-stage iRBC compared to the respective uninfected control. Hierarchical clustering dendrogram was constructed based on the log2FC values of all genes differentially expressed in either of the experimental conditions. **f**, Barplot showing the number of significantly differentially expressed genes (DE genes) (FDR<0.05, log2FC>0.1 or log2FC<-0.1, respectively) in brain endothelial cells. pericytes and astrocytes upon 3D-BBB microvessel exposure to trophozoite and schizont-stage iRBC, or iRBC-egress media compared to the respective uninfected controls.

### The egress of *P. falciparum*-iRBC locally increases barrier permeability and disrupts junctional morphology

Despite the lack of a major transcriptional shift, some of the dysregulated transcripts in 3D-BBB microvessels exposed to trophozoite and schizont-stage iRBC were suggestive of a potential barrier dysfunction. Although TEM did not reveal any changes in the percentage of total junction length covered by tight junctions among the three examined conditions (Fig. 5a,b), we observed junctional differences by immunofluorescence staining. VE-cadherin labelling revealed the presence of thin junctions and inter-endothelial gaps highly localized in microvessel regions near egressed merozoites, which could be identified as small punctate signal (<1µm) by DAPI staining (Fig. 5c). Specifically, the presence of inter-endothelial gaps was minimal in 3D-BBB microvessel regions exposed to high *P. falciparum*-iRBC cytoadhesion with low egress (i.e. where iRBC looked intact and free merozoite were barely present), with 5 (IQR = 4,6) gaps per field of view, compared to 27 (IQR = 19.5, 38) gaps in microvessels with high rate of schizont rupture, largely colocalizing with egressed merozoites (Fig. 5d). Even though no visible changes on the length of tight junctions were observed by TEM, we identified extravasated merozoites in the collagen matrix of 3D-BBB microvessels incubated with schizonts (Fig. 5e). Furthermore, *P. falciparum* schizonts induced functional alterations in barrier integrity, with a 3-fold increase in permeability to 70 kDa FITC-dextran upon 6-hour incubation with schizonts, with median permeability ratios of 0.88 (IQR = 0.51, 1.72) in control and 2.46 (IQR = 1.54, 6.77) in 3D-BBB microvessels exposed to schizonts (Fig. 5f). Collectively, these data suggest that although the effects of *P. falciparum-* iRBC sequestration on endothelial cells are local and associated to the egress of parasite components, they still result into changes in vessel permeability.

**Figure 5.**
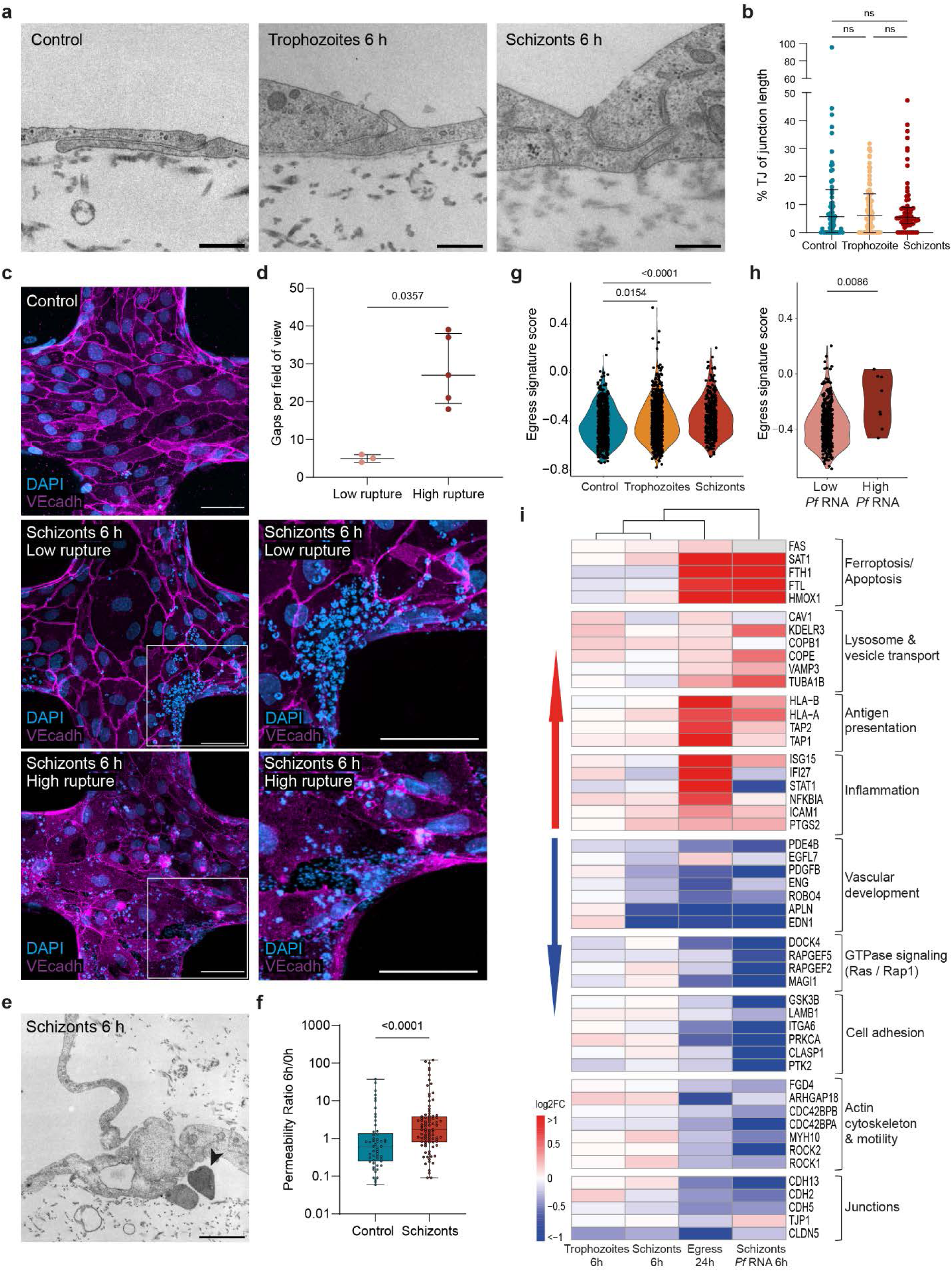
6-hour incubation with *P. falciparum*-iRBC induces localized transcriptional and junctional changes at sites of parasite egress. **a,** Representative TEM images showing intact inter-endothelial junctions formed in 3D-BBB microvessels exposed to trophozoites, schizonts or uninfected RBC. Scale bar = 500 nm. **b**, Scatter plot comparing the percentage of tight junctions per total junctional length upon incubation with control media, trophozoite- and schizont-stage *P. falciparum*-iRBC (Kruskal-Wallis with Dunn’s multiple comparisons test). **c**, Representative maximum z-projection confocal images (left) and close-up views (right) of junctional marker VE-cadherin in a control microvessel and after 6-hour incubation with schizont-stage *P. falciparum*-iRBC. BBB cell and *P. falciparum* nuclei were labeled with DAPI (blue). DAPI staining can be used to identify mature schizonts (∼ 5 μm) and recently egressed merozoites (< 1 μm). Scale bar = 50 μm. **d**, Scatter plot comparing the number of gaps colocalizing with at least one merozoite in microvessels incubated with schizonts at a low (N = 3) or high (N = 5) rupture region. **e**, TEM image showing an extravasated merozoite (arrow head) localized in the collagen hydrogel of 3D-BBB microvessels incubated with schizont-stage *P. falciparum*-iRBC. Scale bar = 2 μm. **f**, Ratio between apparent permeability quantified at 6-hour post-incubation with control media or schizont-stage *P. falciparum*-iRBC and baseline permeability before incubation. Each point represents a different ROI from 3D-BBB microvessel models treated with schizonts (N = 8) or control media (N = 5) Mann-Whitney U test). **g**, Violin plot of *egress signature score* in every endothelial cell plotted by experimental condition (Kruskal Wallis with Dunn’s multiple comparisons test). **h**, Violin plot of *egress signature score* in endothelial cells exposed to schizont-stage iRBC split by *P. falciparum* RNA content (high > 10 logcounts) (*Pf* RNA low: N=385, *Pf* RNA high: N=8) (Mann-Whitney U test). **i**, Heatmap showing log2FC values of selected genes belonging to significant GO-terms shown in Fig. 2f and 3b, in brain endothelial cells after 24-hour exposure to iRBC-egress media, or 6-hour exposure to trophozoite- or schizont-stage iRBC, or the *P. falciparum*-RNA-high endothelial cell population compared to the respective uninfected control. Hierarchical clustering dendrogram was constructed based on the log2FC values of all genes differentially expressed in either of the experimental conditions.

### The egress of *P. falciparum*-iRBC causes localized transcriptional changes

To determine if the local disruption of endothelial cells was accompanied by a transcriptional shift, we defined a *P. falciparum-*iRBC *egress signature score*, including the 50 most upregulated and downregulated genes in endothelial cells exposed to iRBC-egress media (see Methods and Supplementary Information Table 1). Endothelial cells exposed to both trophozoite or schizont *P. falciparum-*iRBC presented a significant increase in the *egress signature score,* which was higher in cells exposed to schizonts (Fig. 5g). Interestingly, the *egress signature score* of endothelial cells exposed to schizont *P. falciparum-*iRBC showed a bimodal distribution compared to the unimodal distribution in the two remaining conditions (Fig. 5g and Extended Data Fig. 5d), suggestive of two transcriptional endothelial states. To infer if this bimodality might be related to spatial proximity to regions of high *P. falciparum-*iRBC egress, we defined an endothelial population that contained *P. falciparum* gene counts above an estimated background-level threshold (see Methods), indicating uptake of parasite material by the respective endothelial cells. Indeed, this population presented a significant increase in the *egress signature score*, compared to cells with a minimal background level of *P. falciparum* reads (p-value = 0.0086) (Fig. 5h). Furthermore, a deeper analysis of the transcriptional alterations in this *P. falciparum*-RNA-high cell population revealed a remarkably similar transcriptional profile to the one of endothelial cells exposed to egress products (Fig. 5i). This included a strong downregulation of transcripts associated with actin cytoskeleton and cell motility, focal adhesions, RAS/RAP1 GTPase pathway, along with low expression of genes encoding for junctional and vascular development pathways. Although only a minor increase in antigen presentation and inflammation pathways was observed compared to iRBC-egress media, an equally strong upregulation of ferroptosis and vesicle transport genes was found (Fig. 5i). Taken together, these results suggest that egress of malaria components from *P. falciparum*-iRBC causes a strong, localized and well-defined signature in endothelial transcription, similar to that found globally in microvessels exposed to *P. falciparum* egress products.

### Pharmacological inhibition of JAK-STAT signaling preserves 3D-BBB integrity

To investigate the relevance of the JAK-STAT pathway as a mediator of the vascular dysfunction caused by *P. falciparum* egress products, and to evaluate its potential as a therapeutic target, we treated our 3D-BBB microvessels with Ruxolitinib, a clinically approved JAK1/2 inhibitor. Briefly, microvessels were incubated for 24 hours with either iRBC-egress media or control cell culture media, in the presence or absence of 10 µM Ruxolitinib. Immunofluorescence staining confirmed that Ruxolitinib prevented the increase in STAT-1 expression induced by *P. falciparum-* egress media (Fig. 6a). Furthermore, the presence of the drug preserved the junctional localization of the tight junction protein ZO-1 in microvessels exposed to *P. falciparum* egress products (Fig. 6b). To determine whether these changes were associated with an improvement in 3D-BBB barrier function, microvascular permeability was assessed by 70 kDa FITC-dextran perfusion before and after treatment, as previously described. While treatment with Ruxolitinib modestly improved baseline permeability in control devices (0.66 – IQR = 0.26, 1.51), its impact was far more pronounced under pathological conditions. Co-incubation with iRBC-egress media led to an almost 24-fold reduction in microvascular permeability (0.45 – IQR = 0.10, 1.94) compared to the iRBC-egress media condition alone (Fig. 6c). Overall, these results highlight the importance of the JAK-STAT pathway in *P. falciparum*-mediated barrier disruption, and support the therapeutic potential of Ruxolitinib in reducing brain vascular dysfunction associated with cerebral malaria.

**Figure 6.**
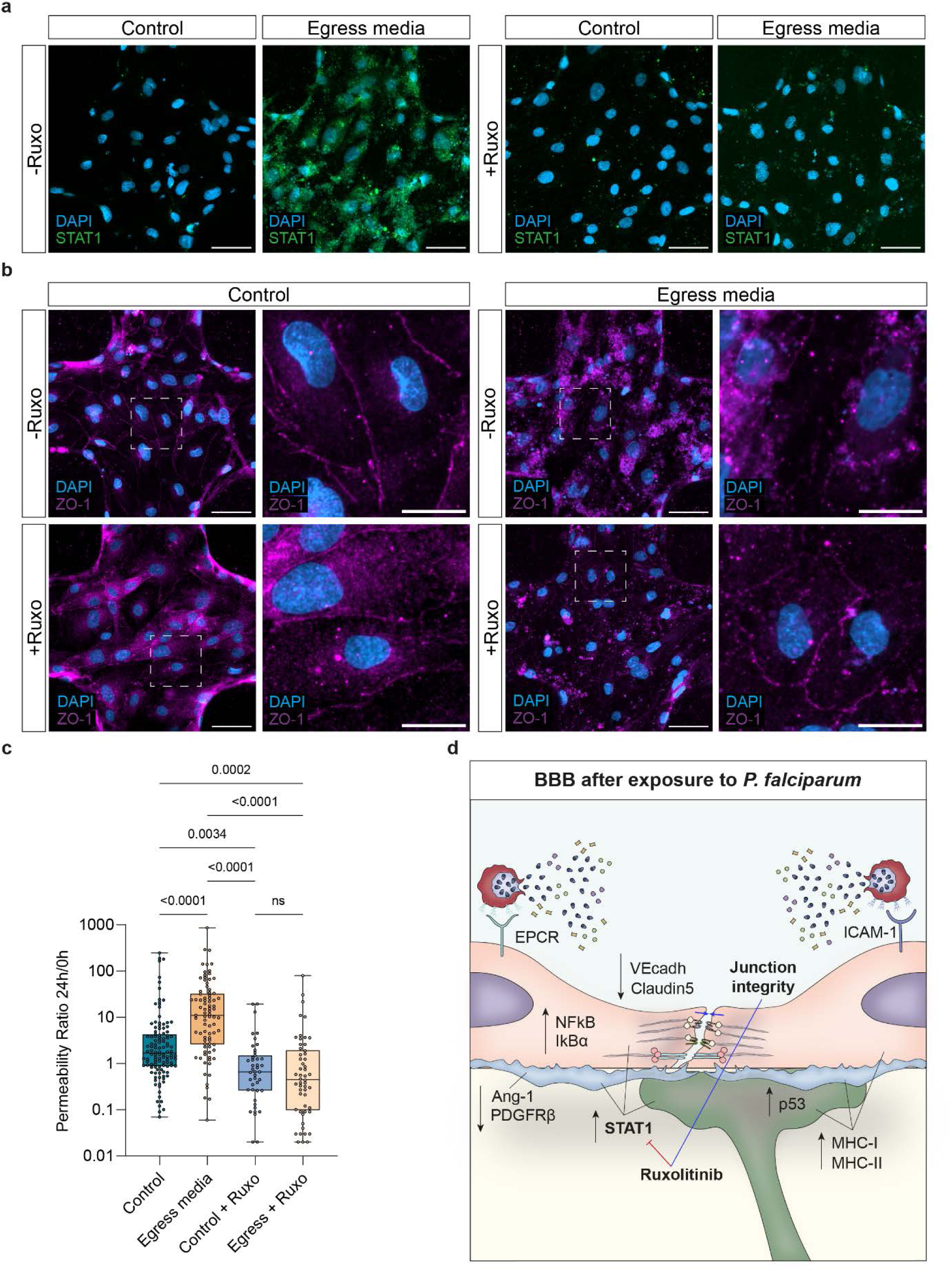
Pharmacological inhibition of JAK-STAT signaling preserves 3D-BBB integrity. **a,** 24-hour co-incubation with Ruxolitinib prevents the overexpression of STAT1 protein in 3D-BBB microvessels, as opposed to the increased expression observed upon exposure to iRBC-egress media alone. Scale bar = 50 μm. **b**, Representative maximum z-projection confocal images (left) and close-up views (right) of junctional marker ZO-1 (magenta) in control microvessels and after 24-hour incubation with iRBC-egress media, and upon co-incubation with Ruxolitinib. BBB cell nuclei and *P. falciparum* DNA were labeled with DAPI (blue). Scale bar = 50 μm; close-up scale bar = 20 μm. **c**, Ratio of the apparent permeability calculated at 24-hour post-incubation and at pre-incubation with control media or iRBC-egress media. Each point represents the ratio from an ROI coming from 3D-BBB microvessels exposed to control media (N = 4), iRBC-egress media (N = 10) or co-incubated with Ruxolitinib and control media (N = 3) or iRBC-egress media (N = 4) (Kruskal Wallis with Dunn’s multiple comparisons test). **d**, Schematic representation of the altered pathways in the BBB cell types upon iRBC-egress media exposure and at sites of natural iRBC rupture. *P. falciparum* egress products released upon iRBC rupture activate antigen presentation (MHC I, MHC II) and inflammation-related (STAT1) pathways in BBB cell types. Furthermore, while endothelial cells downregulate junctional markers, pericytes show reduced Ang-1 and PDGFRβ signaling and astrocytes activate the p53 apoptosis-associated pathway. Pharmacological inhibition of JAK-STAT pathway through Ruxolitinib prevents STAT1 overexpression and preserves junction integrity.

## Discussion

CM is characterized by *P. falciparum* accumulation in the brain microvasculature and it is often accompanied by vascular dysfunction^2,3^. Despite the severity of the disease, our understanding of how the malaria parasite affects the BBB remains limited, primarily due to difficulties in obtaining brain samples from affected patients or the lack of accurate disease models. In this study, we developed a bioengineered microvascular 3D-BBB model that incorporates primary human brain microvascular endothelial cells, pericytes and astrocytes. The addition of these cell types increased the vascular barrier function of the model, improving upon other bioengineered models previously used to study *P. falciparum* pathogenesis^24,30,31^. We used this advanced model to assess BBB disruptive mechanisms mediated by *P. falciparum.* Perfusion and incubation either with media containing *P. falciparum-*egress products or with *P. falciparum* schizonts led to a global increase in microvascular permeability to 70 kDa FITC-dextran. However, differences in the extent of vascular barrier opening were found between these two conditions. While large VE-cadherin inter-endothelial gaps formed after incubation with iRBC-egress media, schizont sequestration only induced smaller gaps near areas of high merozoite egress. Similar differences in the extent and ubiquity of parasite-induced transcriptional shifts were found in our scRNA-seq analysis. We observed major and widespread alterations in endothelial gene expression after exposure to iRBC-egress media, and only minimal global differential expression after 6-hour incubation with trophozoites and schizonts. This analysis is in agreement with other bulk transcriptomics studies that showed limited endothelial transcriptional changes upon incubation with *P. falciparum-* iRBC^30^ and more prominent changes after incubation with *P. falciparum* lysates^29^. Nevertheless, the single cell resolution of our transcriptomic analysis revealed that endothelial cells near regions of egress, identified by high *P. falciparum* RNA content, presented a transcriptional signature similar to that of endothelial cells exposed to iRBC-egress media. Altogether, our results show that although *P. falciparum-*mediated disruption is highly localized to areas of *P. falciparum* egress, it could still result in severe pathogenic outcomes, as shown by the increase in barrier permeability we observed.

Although *P. falciparum* has a broad repertoire of members of the parasite ligand PfEMP1, CM patients are enriched in variants that bind to EPCR^41^. Whether parasite binding to specific endothelial receptors directly contributes to vascular disruption remains unknown. Recent studies have reported conflicting results on the transcriptional effect of cytoadherent *P. falciparum-*iRBC. Studies on endothelial-only 3D brain microvessels have shown minimal transcriptional differences after short term incubation with *P. falciparum*-iRBC^30^. Likewise, no differences in key endothelial transcripts were found between parasites derived from CM and uncomplicated malaria patients in a study in Malawi^42^. Conversely, a recent study has shown differential endothelial gene expression upon binding of parasite lines expressing different PfEMP1^43^. Our study did not find major transcriptional differences after short term incubation with highly synchronized trophozoites expressing a dual EPCR-ICAM-1 binding PfEMP1 previously shown to be highly disruptive in *in vitro* BBB spheroids^14^. Overall, the lack of transcriptional differences could be a result of the short incubation period, chosen to disentangle the effects of binding from that of natural parasite egress. We cannot rule out the possibility of a *P. falciparum* binding-induced transcriptional shift at later time points, and the occurrence of non-transcriptional cellular processes not evaluated in our study, including morphological or mechanical modifications on endothelial cells, such as transmigratory-like cup structures or trogocytosis, previously identified in *P. falciparum in vitro* studies^44^. Other important disruptive effects not investigated in this study are synergistic damaging effects in the presence of other co-factors, such as protein C and thrombin^30,45^.

Our study confirms that the egress of sequestered *P. falciparum* parasites in close proximity to endothelial cells is a key pathogenic event, accompanied by the release of agents such as heme^17^, hemozoin^18^, parasite histones^19,20^, PfHRP2^21^, and GPI anchors^23^, which have been previously described to activate endothelial cells and compromise barrier integrity. We reproduced *P. falciparum-*induced transcriptomic signatures found in other studies^29,30^, including those related to endothelial disruption of ER-transport, oxidative stress and ferroptosis. These changes likely resulted from the detoxification of hemozoin, heme and other parasite products^46^. Nevertheless, the use of an improved 3D-BBB model revealed new parasite disruptive mechanisms. We showed for the first time that *P. falciparum* induces a global endothelial downregulation of the tight junction marker *CLDN5* (Claudin-5) in all the parasite conditions we tested, as well as the decrease of other junctional transcripts in microvessels treated with iRBC-egress media (Fig. 6d). Of additional relevance is the decrease in expression of vascular developmental genes, as well as in transcripts from pathways related to cell adhesion and cytoskeletal organization. Our study also shows the downregulation of important homeostatic signaling pathways between endothelial cells and pericytes^47^, including the PDGF-PDGFR^48^ and Notch signaling pathways^49^, and the Angiopoietin/Tie-2 axis^31^. These data are in agreement with a recent study from our group demonstrating that pericytes play a functional role in CM pathogenesis by halting their expression of Ang-1^31^. Upregulated pathways in astrocytes and pericytes in response to iRBC-egress media include collagen secretion, suggestive of fibrosis, and p53-mediated astrocytic apoptosis, which could explain the reduced astrocyte density observed by immunofluorescence staining. However, we did not observe any increase in astrocyte activation and GFAP expression in response to *P. falciparum.* Post-mortem studies on CM samples have revealed different degrees of astrogliosis, although not co-localizing with iRBC sequestration sites^3^, suggesting that astrocyte reactivity may be caused by a different pathogenic driver.

Our study has revealed that all the cell types included in the 3D-BBB model present upregulation of type I IFN response and antigen presentation pathways. Interestingly, polymorphisms of the IFN-alpha receptor-1 (*IFNAR1*)^50^ or *IFIT1*^51^ have been associated with a reduced risk of CM. Our findings align with previous reports of type I IFN response in a *P. berghei* experimental cerebral malaria model^52^. Other studies on *P. berghei* CM models have suggested a potential mechanism of merozoite engulfment and cross-presentation of parasite antigens by endothelial cells ^34,35^, or by astrocytes and microglia^53^. Our model suggests the existence of similar engulfment mechanisms of *P. falciparum* egress products, likely responsible for the activation of antigen presentation pathways in endothelial cells, but also in pericytes and astrocytes within the collagen hydrogel. Our findings and others^44,52,53^ suggest a potential mechanism for leukocyte recruitment not only intravascularly by endothelial cells^30^, but also at the brain perivascular space, aligning with recent observations of vascular and perivascular accumulation of CD8+ T-cells in the brain microvasculature in post-mortem samples of CM patients^54,55^. In this scenario, our results suggest that blood-brain barrier cells might be able to present internalized parasite material to CD8+ T-cells, which could potentially amplify damage at the vascular and perivascular space.

Another inflammation-associated response highly upregulated in the three BBB cell types was the JAK-STAT pathway, as indicated by the strong transcriptional activation of members of this pathway, as well as by the nuclear translocation of STAT1 in endothelial cells after incubation with iRBC-egress media. Enhancement of leukocyte activation of type I and II IFN and pro-inflammatory pathways, such as JAK-STAT, is a prominent systemic feature in *P. falciparum* infection (reviewed in ^56^). Notably, to explore the therapeutic potential of targeting this pathway, we treated 3D-BBB microvessels with Ruxolitinib, a clinically approved JAK1/2 inhibitor. Our data demonstrate that Ruxolitinib effectively prevented barrier disruption induced by *P. falciparum* egress products, as evidenced by preserved tight junction integrity and reduced microvascular permeability (Fig. 6d). Interestingly, Ruxolitinib also conferred a moderate protective effect in the absence of *P. falciparum*-associated stimuli, suggesting the dual capacity of JAK-STAT inhibition to stabilize endothelial function both under inflammatory and resting conditions. Recent studies in clinical human malaria infections have shown the tolerability of Ruxolitinib in combination with antimalarials^57^, and its efficacy as an adjunctive therapy in uncomplicated infections by attenuating inflammation and endothelial activation early in infection^58^, without affecting immune memory responses^58,59^. Altogether, our results, combined with emerging data in human infected volunteers, suggest that inhibitors of the JAK-STAT pathway, such as Ruxolitinib, could be repurposed not only to modulate systemic immune responses but also to directly protect vascular integrity and mitigating cerebral complications during *P. falciparum* infection.

While our 3D-BBB model represents a significant advancement over previous *in vitro* systems, it still presents several limitations. Microvessel fabrication is complex and requires a long period of training in experienced laboratories. Although presenting improved barrier properties, our model is still far away from recreating BBB physiological permeability rates. While the barrier properties of our 3D-BBB microvessels could be enhanced using induced pluripotent stem cell (iPSC)-derived brain endothelial cells^60^, concerns have been raised about their epithelial-like characteristics^61^, particularly in early protocols. Recent advances have shown that overexpression of ETS factors, such as ETV2, FLI1, ERG ^61^, or FOXF2 and ZIC3^62^ can shift these cells towards an improved vascular and brain vascular phenotype, respectively. As other differentiation mechanisms are currently being developed for the generation of iPSC-derived BBB models^63,64^, iPSCs still remain a promising tool for generating *in vitro* models of the microvasculature of the brain and other organs affected during *P. falciparum* infection^65^. Another limitation of our study is that we have solely focused on the pathogenic effect of *P. falciparum* and have not evaluated the effect of cytokines^39^ or immune cell types^54,66^. Future studies could take advantage of the controlled microfluidic properties of our model to introduce these components, which play a relevant role in CM. Finally, although our 3D-BBB model presents increased complexity, other brain cell types, including microglia and neurons, could be incorporated to further resemble the human physiology and investigate brain-associated pathogenic mechanisms. Despite these limitations, our model has shed light on novel pathogenic pathways of *P. falciparum* malaria and highlights the value of using innovative bioengineered models to enhance our understanding of infection and facilitate the development of future treatments.

## Methods

### Primary human cell culture

Primary human brain microvascular endothelial cells (HBMEC, Lot 376.05.04.01.2F or 376.11.04.01.2F, Cell Systems) were cultured in endothelial cell growth medium-microvascular (EGM-2MV, Lonza) up to passage 8. Primary human astrocytes (HA, Lot 31978, ScienCell) were cultured in basal media supplemented with 2% FBS, 1% Pen-Strep solution and 1% astrocyte growth supplement (ScienCell) up to passage 6. Primary human brain vascular pericytes (HBVP, Lot 32562, ScienCell) were cultured in basal media supplemented with 2% FBS, 1% Pen-Strep solution and 1% pericyte growth supplement (ScienCell) up to passage 8. All cell types were cultured at 37°C and 5% CO2 as monolayers until microvessel fabrication, using a poly-L-lysine (0.1% (w/v), Sigma-Aldrich) coating for cell attachment.

### Green fluorescent protein and mCherry lentiviral transduction

GFP-expressing HA and mCherry-expressing HBVP were obtained by lentiviral transduction. Briefly, cells were grown in a T75 flask and, once confluent, they were incubated for 24 hours in serum-free astrocyte or pericyte media with concentrated viral particles containing a GFP or mCherry vector (kindly donated by the laboratory of Dr Kristina Haase, EMBL Barcelona), at a multiplicity of infection of 10. After 24 hours, the lentiviral particles were removed with two consecutive 24-hour washes in their respective cell culture media before astrocytes and pericytes were expanded and frozen down. For GPF-expressing HA, the efficiency of transduction was quantified as percentage of GFP signal overlapping with cell-covered area. mCherry-positive pericytes were further selected by fluorescence-activated cell sorting.

### P. falciparum culture

HB3var03 *P. falciparum* parasites were cultured using human 0+ erythrocytes in RPMI 1640 medium (GIBCO) supplemented with 10% human type-AB+ plasma, 5mM glucose, 0.4 mM hypoxantine, 26.8 mM sodium bicarbonate and 1.5 g/L gentamicin (RPMI complete). Parasites were grown in a gas mixture of 90% N2, 5% CO2, and 5% O2 and parasitemia was regularly checked by Giemsa staining to avoid culture overgrowth. Cultures were regularly panned and monitored for correct PfEMP1 expression. *P. falciparum* parasite were synchronized weekly using 5% sorbitol to select for ring-stage parasites and 70% Percoll gradient to select for schizonts.

### 3D-BBB microvessel fabrication

The protocol for 3D microvessel fabrication can be found in ^25^, and we conducted the following modifications for the generation of a 3D-BBB model. Briefly, type I collagen was isolated from rat tails as previously described and dissolved in 0.1% acetic acid to a stock concentration of 15 mg/mL before dilution to 7.5 mg/mL and neutralized on EGM-2MV supplemented with 1% astrocyte and pericyte growth factors (ScienCell) before HBMEC seeding for microvessel fabrication. Primary HA and HBVP were added to the neutralized collagen solution in a 7:3 ratio, using a concentration of 7.5x10^5^ HA/mL(collagen): 3.2x10^5^ HBVP/mL(collagen). A 1:1 astrocyte-to-pericyte ratio (2.5x10^5^ HA/mL(collagen): 2.5x10^5^ HBVP/mL(collagen)) was initially used to compare the expression level of BBB-specific markers by quantitative polymerase chain reaction. A multi-step process combining soft lithography and injection molding was used to fabricate microvessels, as previously reported^25,67^. Briefly, the top and bottom parts of the microvessels were fabricated separately and then assembled within two polymeric housing jigs. The negative impression of a microfluidic network was obtained by injecting the collagen solution between the top polymeric jig and a positive polydimethylsiloxane (PDMS) micro-patterned mold, previously made hydrophilic by O2 plasma treatment. The bottom part was fabricated by pouring the collagen solution on top of a 22 mm x 22 mm coverslip within the bottom housing jig and by compressing it using a flat PDMS stamp to obtain a thin collagen layer. The two pieces were left gelling up to 1 hour at 37°C and then assembled after removal of the PDMS molds. The microvessels were incubated for at least one hour with EGM-2MV medium supplemented with 1% astrocyte and pericyte growth factors (ScienCell) before HBMEC seeding. Primary HBMEC were seeded at a concentration of 7x10^6^ cells/mL under gravity-driven flow by adding 8 µL volume increments to the device inlet until reaching full coverage in the microfluidic network. Microvessels were cultured for up to 7 days and fed every 12 hours by gravity-driven flow.

### Quantitative polymerase chain reaction

After microvessels fixation at 0.5, 3 or 7 days in culture, total RNA was isolated from disassembled collagen hydrogels using TRIzol and then purified by RNeasy Mini Kit (Qiagen 50974004). The purified RNA was quantified using a NanoDrop™ 2000c Spectrophotometers (ThermoFisher) and then converted to the complementary cDNA using the TaqMan Reverse Transcription Reagents (ThermoFisher, N8080234) according to the manufacturer’s instructions. Quantitative polymerase chain reaction (qPCR) was performed using the LightCycler 480 SYBR Green I Master (Roche, 04707516001) in a LightCycler 480 II (Roche). The oligonucleotides used as primer sequences for the qPCR experiments were purchased from ThermoFisher and are reported in Extended Data Table 1. The PCR program consists of an initial step at 95°C for 15 minutes, followed by 45 cycles of 30-second denaturation at 94°C and 40-second annealing at 60°C and 50-second extension at 72°C. The automatically detected threshold cycle values were compared using the ΔΔCt method, with the 0.5-day condition as the reference for comparison, and the gene expression levels were normalized to those of the housekeeping gene PECAM1.

### Immunofluorescent staining of 2D monolayers

HBMEC, HA or HBVP were seeded on poly-L-lysine-coated 8-well slides (Falcon) at a concentration of 2x10^4^ cells/well and grown until reaching confluency. 2D monolayers were fixed for 20 minutes with ice-cold 4% PFA. Fixation was followed by two consecutive phosphate-buffered saline (PBS) washes and a 1-hour blocking-permeabilization solution in 2% BSA and 0.1% Triton X-100 in PBS at room temperature. Primary antibodies were diluted in a 2% BSA and 0.1% Triton X-100 PBS solution and incubated for 1 hour at room temperature. Primary antibodies against the following proteins were used: VE-cadherin (Santa Cruz Biotechnology sc-52751 or Abcam ab33168), STAT1 (Cell Signaling 14994S), vWF (Bio-Rad AHP062), PECAM1 (BD Pharmingen 560983), β-catenin (Santa Cruz Biotechnology sc-59737), ICAM-1 (Abcam ab20), GFAP (Abcam ab4674), S100B (Sigma S2532-100U), AQP4 (Novus Biologicals NBP1-87679), αSMA (Abcam ab202509), PDGFRβ (Abcam ab69506), NG2 (Invitrogen 372700). After two PBS washes, the monolayers were incubated with 2 µg/mL DAPI (ThermoFisher D21490, 1:250), Alexa-Fluor 488-, Alexa-Fluor 594- or Alexa-Fluor 647-conjugated secondary antibodies (Invitrogen, 1:250) for 1 h at room temperature and then washed twice with PBS. Images were acquired using a LSM980 Airyscan 2 microscope (Zeiss) and processed with imaging software ZEN (Zeiss) and Fiji (ImageJ).

### Immunofluorescent staining of 3D microvessels

3D-BBB microvessels were fixed with ice-cold 4% paraformaldehyde (PFA) for 20 minutes after 7 days in culture. All the solutions for 3D microvessel staining were perfused through gravity-driven flow. Fixation was followed by two consecutive 10-minute PBS washes before immunofluorescent staining. Any possible background signal coming from the collagen hydrogel was quenched using Background Buster (Innovex Biosciences) for 30 minutes. After blocking/permeabilization in 2% BSA and 0.1% Triton X-100 (in PBS) at room temperature for 1 hour, the devices were incubated at 4 °C overnight in a PBS solution containing the primary antibodies, 2% BSA and 0.1% Triton X-100. Primary antibodies against the following proteins were used: GFP (Invitrogen A21311, 1:200), mCherry (Invitrogen M11240, 1:200), VE-cadherin (Santa Cruz Biotechnology sc-52751, 1:100 or Abcam ab33168, 1:100), STAT1 (Cell Signaling 14994S, 1:100), αSMA (Abcam ab202509, 1:200), LAMP1 (Cell Signaling 9091, 1:100), ZO-1 (Invitrogen 339100, 1:100). After six 10-minute PBS washes, the microvessels were incubated with 2 µg/mL DAPI (ThermoFisher D21490, 1:250), Alexa-Fluor 488-, Alexa-Fluor 594- or Alexa-Fluor 647-conjugated secondary antibodies (Invitrogen, 1:250) for 1 h at room temperature and then washed six times for 10 minutes with PBS. Imaging of the devices was performed using a LSM980 Airyscan 2 microscope (Zeiss) and images were processed with imaging software ZEN (Zeiss), Fiji (ImageJ) and Vision4D (Arivis)

### Confocal image analysis

Confocal images were analyzed using Fiji (ImageJ) for quantification of different parameters. For each image, Z-stack slices were summed to preserve all the fluorescent signal and the images were split into different channels and analyzed as 2D images. Quantification of gaps at sites of *P. falciparum*-iRBC rupture was obtained manually counting the number of gaps that colocalized with at least one free merozoite. Gap number per field of view was expressed as mean ± standard deviation. For nuclear STAT1 quantification, the DAPI channel was used to create a mask of the nuclei to select the regions of interest (ROIs). For each ROI, we measured the mean fluorescent value in the STAT1 channel and we then compared the median nuclear fluorescent intensity among conditions. To obtain astrocyte density, the DAPI channel was used to create a mask and count the nuclei per field of view.

### P. falciparum-iRBC egress media preparation

The protocol for generation and purification of *P. falciparum*-iRBC can be found in^31^. Briefly*, P. falciparum*-iRBC at 42-48 hours post-infection (hpi) were purified by a gelaspan (40 mg/mL) gradient separation and treated for 5 hours with the reversible PKG inhibitor C2 (kindly donated by Michael Blackman, The Francis Crick Institute) in RPMI complete media to inhibit parasite egress. After drug removal, parasites were resuspended at a concentration of 10^8^ *P. falciparum*- iRBC/mL in EGM-2MV media supplemented with 1% astrocyte and pericyte growth factors (ScienCell), gassed with a mixture of 90% N2, 5% CO2, and 5% O2 and left in the incubator overnight on a shaker (50 rpm) to facilitate parasite egress. The efficiency of parasite egress was assessed by hemocytometer count and blood smear. The concentration was adjusted at 5x10^7^ ruptured *P. falciparum*-iRBC/mL before the suspension was centrifuged at 1000 rpm for 5 minutes, aliquoted and flash frozen in liquid nitrogen. The same protocol was applied to uninfected erythrocytes to be used as a negative control medium for the scRNA-seq experiment.

### Sample incubation with P. falciparum egress media, Ruxolitinib, P. falciparum-iRBC or cytokines

After 6 days in culture, 3D-BBB microvessels were perfused with 150 µL of *P. falciparum*-iRBC egress media under gravity-driven flow and incubated for 24 hours at 37°C. Reservoirs were refilled every 12 hours. For Ruxolitinib (Santa Cruz Biotechnology) treatment, 3D-BBB microvessels were co-incubated for 24 hours at 37°C using a 10 µM Ruxolitinib concentration and iRBC-egress media or cell culture media.

For trophozoite and schizont *P. falciparum*-iRBC perfusion, a magnetic cell separation or Percoll gradient was used to purify late-stage parasites at the desired hpi (26-34 hpi for trophozoites or 42-48 hpi for schizonts) at >60% purity. Microvessels were then perfused thrice for 10 minutes with 150 µL of *P. falciparum*-iRBC at 5x10^7^ iRBC/mL concentration under gravity-driven flow followed by two consecutive 10-minute washes to remove unbound cells. Devices were then incubated at 37°C with *P. falciparum*-iRBC for 6 hours. As a control, the same concentration of uninfected erythrocytes or uRBC media was used for perfusion and incubation. After incubation, samples were used for single cell RNA sequencing, permeability assays or immunofluorescence staining for confocal imaging with the experimental timeline shown in Fig 2a and 4a.

Confluent 2D cell monolayers in 8-well slides were incubated with 150 µL of *P. falciparum*-iRBC egress media (50x10^6^ ruptured iRBC/mL). Cell culture media was used as control. Alternatively, 2D monolayers were incubated overnight with TNFα (R&D Systems), IL-1β (Peprotech) and interferon γ (IFNγ) (Peprotech), either alone or combined in a cytokine cocktail. All cytokines were used at a concentration of 10 ng/mL. Cell culture media was used as control.

### Sample preparation and imaging by transmission electron microscopy

Microvessels were pre-fixed for 30 min in 2% PFA / 2.5% glutaraldehyde (GA) in EGM-2MV medium for 30 minutes and washed thrice for 10 minutes with EGM-2MV. The collagen hydrogel was carefully removed from the PMMA jig and the low-shear stress areas of the microvessel network were cut into smaller pieces (about 1 x 0.5 x 0.5 mm) for further processing. The samples were fixed with a secondary fixative solution (2% PFA, 2.5% GA, 0.25 mM CaCl_2_, 0.5 mM MgCl and 5% sucrose in 0.1 M sodium cacodylate buffer) overnight at 4°C, rinsed twice for 15 minutes with 0.1 M sodium cacodylate buffer and stained with reduced osmium solution (1% OsO_4_, 1.5% K_3_FeCN_6_ in 0.065 M Cacodylate buffer) for 2 hours at 4°C. Samples were washed six times for 10 minutes in distilled H_2_O and kept at 4°C until further processing. Dehydration was performed in steps of 30, 50, 80, and thrice with 100% ethanol in a PELCO Biowave Pro microwave processor (Ted Pella, Inc.) containing a SteadyTemp Pro and a ColdSpot set to 4°C, each step 40 seconds at 250 W. Samples were infiltrated in serial steps of 25, 50, 75, 90, and twice with 100% EPON 812 hard epoxy resin in acetone, assisted by the microwave (3 minutes each at 150 W under vacuum) and a final infiltration step in 100% EPON 812 hard epoxy resin overnight at room temperature. Samples were oriented in the embedding mold with the axis of the microvessel lumen at approximately 90° angle to the cutting surface to be able to cut transversal sections of the channels, and then polymerized at 60°C for 48 hours. Microvessel pieces for imaging were randomly selected and thin sections (70 nm) were retrieved on an ultramicrotome (UC7, Leica Microsystems), collected on formvar-coated slot grids and post-stained in uranyl acetate and lead citrate. Tile montages (12,000x) were acquired on a JEOL JEM 2100 plus at 80 or 120 keV using SerialEM. Montages were processed using IMOD’s Blend Montages function and Fiji (ImageJ). Tight junctions were analysed as percentage of length of electron-dense tight junctions over junction length as measured in Fiji (ImageJ). Percentage of gaps in junctions was counted in Fiji (ImageJ).

### Microvascular permeability assays

Permeability assays were performed on endothelial-only or 3D-BBB microvessel models after 6 and 7 days in culture. 70 kDa FITC-dextran at a concentration of 100 µg/mL was perfused at a flow rate of 10 µL/min, applied with a withdrawing syringe pump (Harvard Apparatus PHD 2000). Tilescan confocal images were acquired with 5 μm z-step size in 5 different ROIs of the microvessels every 2.5 minutes over 10 minutes. For time lapse experiments, imaging was performed on the same ROI before and after 24 hour-incubation with *P. falciparum* egress media or 6-hour incubation with schizont-stage *P. falciparum*-iRBC.

Microvascular permeability quantification. The quantification of apparent microvascular permeability was done using the following formula:

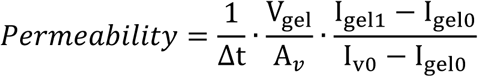

The leakage of the fluorescent tracer into the collagen matrix between two time points was calculated as follows, were Δt is the time interval between the two frames, V_gel_ is the volume of the collagen matrix, A_v_ is the lateral vessel surface, I_gel1_ – I_gel0_ is the difference between the fluorescence intensity inside the gel in the two time points, I_v0_ is the fluorescence intensity in vessel at the start of the measurement. An area of determined size (250 µm x 150 µm) containing part of the vessel and the collagen matrix was selected after image analysis in Fiji (ImageJ). Two ROI were defined, corresponding to the vessel (250 µm x 30 µm) and the adjacent collagen matrix (250 µm x 120 µm) (Extended Data Figure 1). For each ROI, fluorescence intensities were obtained at two different time points -t0 (after complete vessel filling with dextran) and t1 (2.5 minutes later). These values were then used to calculate microvascular permeability (cm/s). Permeability was calculated before incubation with iRBC-egress media or schizonts iRBC, and after 24 or 6 hours, respectively. A final/baseline permeability ratio was calculated for each microvessel area and used to compare different conditions.

### Statistical analysis

Statistical analysis was performed using GraphPad Prism (version 10.2.0) or R (version 4.2.2). Two-tailed Mann-Whitney U test was used to analyze non-normally distributed samples. The effect size *r* was calculated as the *z*-statistic divided by the square root of the sample size *n*. To compare non-normally distributed samples among multiple conditions, Kruskal-Wallis test with Dunn’s multiple comparisons test was used. P values < 0.05 were considered statistically significant. Values are reported as median (Interquartile Interval), each dot in the graphs represents a ROI or an event from a number of replicates (N) indicated in the figure legends.

### Sample preparation for single-cell RNA sequencing (scRNA-seq)

The 3D-BBB models were fabricated, perfused and incubated according to the described conditions. 3-4 devices per condition were disassembled, and the main, cell-containing collagen part was dissected. The collagen piece was dissociated for 6-10 minutes (until complete dissociation) in collagenase diluted in serum-free media (1 mg/mL, Sigma-Aldrich). After stopping the collagenase reaction with complete media, the cells were trypsinized for 8 minutes to obtain a single-cell solution. The cells were mechanically dissociated by pipetting with wide-bore tips and washed two times in PBS containing 0.1% BSA.

### MULTI-seq sample preparation

The single-cell solutions harvested from the 3D-BBB microvessels were labeled with MULTI-seq barcode oligonucleotides for sample multiplexing as described^32^. Briefly, the cells were resuspended in Cell Prep Buffer (PBS containing 0.1% poly(vinyl alcohol) (PVA) and 1 mM EDTA). A 1:1 mixture of the cholesterol-conjugated Anchor-oligonucleotides (Anchor CMO, synthesized by Integrated DNA Technologies) and Barcode oligonucleotides with a distinct barcode for each sample (final concentration, 0.2 µM) was added and the cells were incubated on ice for 5 min. Next, the same concentration of Co-Anchor CMO (synthesized by Integrated DNA Technologies) was added and incubated for another 5 min, followed by three washes with PBS containing 1% BSA. The cells in each sample were counted after washing and were combined so that the multiplexed suspension contained the same numbers of cells from each sample. The combined sample was filtered through a 35 µm cell strainer and counted again before 10x Genomics barcoding. The MULTI-seq barcode sequences used in this study are: *TCCTCGAA* for control uRBC media, *ATGCGATG* for iRBC-egress media, *GCTATGCA* for control RBC, *CGATACTG* for trophozoite stage iRBC, *TACGCAGT* for schizont stage iRBC.

*10x Genomics barcoding and sequencing.* mRNA transcripts of each cell were barcoded using the Chromium Controller (10x Genomics, firmware version 4.00). The reagent system was Chromium Single Cell 3′ GEM, Library & Gel Bead Kit v3.1 (10x Genomics) and a Chromium Next GEM Chip G Single Cell Kit (10x Genomics). Barcoding and cDNA library construction were performed according to the manufacturer’s instructions and MULTI-seq barcode library preparation was performed as per the MULTI-seq protocol^32^. After the cDNA amplification step, the barcode fraction was collected, amplified, and single-indexed with KAPA HiFi HotStart ReadyMix (Roche) for sequencing. Both finished cDNA and MULTI-seq-barcode libraries were sequenced with NextSeq2000 (Illumina). We read 8 base pairs (bp) for TruSeq Indices, 28 bp for 10x Genomics barcodes and unique molecular identifiers (UMIs), and 52 bp for both fragmented cDNA and MULTI-seq barcodes.

### scRNA-seq data analysis

*Sequence alignment.* Sequenced reads were aligned to a combined reference genome constructed from the human genome (*GRCh38*) and the *P. falciparum* genome (*hb3*) to generate the feature-barcode matrices with the *CellRanger* pipeline (v. 7.0.1, 10x Genomics). All downstream data analysis was performed with *R version 4.2* ^68^.

*Demultiplexing of the MULTI-seq sample.* Using the R package *deMULTIplex* (v.1.0.2)^32^ we counted the MULTI-seq barcode reads and assigned each cell to being a singlet of a specific condition, a doublet, or a sample-barcode-negative cell. Only singlets were kept for further analysis.

*Quality Control (QC).* We performed quality control using the *scuttle* package (v.1.8.4)^69^ as follows: Genes that were found in less than 5 cells were excluded. We then identified the QC thresholds based on scatter plots of detected gene counts against the proportion of mitochondrial gene expression in each cell or the *isOutlier* function (nmads=1.5) and only kept the cells above the determined detected gene cutoff (iRBC-egress dataset: 2500, iRBC dataset: 3233/2448) and below the mitochondrial gene cutoff (iRBC-egress dataset: 4%, iRBC dataset: 5%) for further analysis. Subsequently, doublets were excluded using the *scDblFinder* package (v. 1.10.0)^70^.

*Data normalization.* We normalized the raw UMI counts of the QC-filtered cells using the deconvolution approach from the R package *scran* (v.1.24.1)^71^. The size factor for the library size correction of each cell was calculated with the *calculateSumFactors* function and the raw counts were normalized based on the size factor and log2-transformed with the *logNormCounts* function. These values appear as “*logcounts*”.

*Identification of highly variable genes (HVG), visualization, and clustering.* After normalization, highly variable genes were chosen with the *modelGeneVar* function in the scran package and genes with a biological variance > 0.1 were chosen as HVGs for further dimension reduction and data integration. Using the chosen HVGs we performed principal component analysis (PCA) and constructed UMAP plots from the principal components capturing the highest percentage of variance using the *scater* package (v.1.26.1)^69^. The gene expression levels shown in the UMAP plots correspond to the log2-transformed normalized values. Cell population clusters were identified using the *bluster* package (v.1.8.0)^72^ and the *Leiden* algorithm^73^. Cell type assignment was performed manually based on the cluster marker genes.

*Data integration.* The dataset of the trophozoite-/ schizont-iRBC perfusion was the only dataset where two 10x reactions were used while both reactions contained all three MULTI-seq-labeled conditions (RBC-/ trophozoite-/ schizont-iRBC perfusion). To integrate the two sequencing runs the data was scaled according to sequencing depth using the multiBatchNorm function in the R package *batchelor* (v.1.14.1)^74^. HVGs across the datasets were chosen using the *combineVar* function. The *rescaleBatches* function was used for data integration and the corrected results were used for all visualizations, but not for differential gene expression analysis.

*Differential expression analysis.* Differential expression analysis was performed separately for each cell type utilizing the hurdle (two-part generalized regression) model from the *MAST* package (v.1.22.0)^75^. The Benjamini-Hochberg method was applied to the p-values to account for multiple testing. All significantly differentially expressed genes for all cell types and conditions can be found in Supplementary Information Table 1. The heatmaps visualizing the log2-transformed fold change (log2FC) values were assembled using the *pheatmap* (v.1.0.12)^76^ and the *dendextend* package (v.1.16.0)^77^. The heatmap genes were selected as representatives of the most significantly overrepresented GO-term from the endothelial differentially expressed genes upon iRBC-egress exposure.

*Gene ontology (GO) term over-representation analysis.* GO-term over-representation analysis on significantly up- or downregulated genes was performed separately per each cell type and condition (FDR<0.05, log2FC > 0.1 or < -0.1, respectively)) using the *enrichGO* function (Benjamini-Hochberg correction, p-value cutoff 0.05, max. geneset size 1500) from the *clusterprofiler* package (v.4.4.4)^78^. The iRBC-egress dataset included “*Biological Process*” and the iRBC dataset included all GO-term categories. The *pairwise_termsim* and *emapplot* functions from the *GOSemSim* package (v.2.22.0)^79^ and the *enrichplot* package (v.1.16.2)^80^ were used to visualize the analysis results. GO-term clusters in the enrichment map were manually labeled. The list of all significant GO-terms can be found in Supplementary Information Table 2.

*Gene signature analysis.* Gene signature scores were calculated using the *tidySingleCellExperiment* package (v.1.6.3)^81^. Log-expression values of all signature genes were summed up and a signature score was calculated by rescaling the resulting number to a signature score between 0 and 1 for every single cell:

*MHC signature score = rescale(sum(gene log-expression values), to=c(0,1))*

The genes included in the MHC signature scores were obtained from the KEGG pathway^82^ “*hsa04612-Antigen processing and presentation*”. These include

MHC1: *PSME3, PDIA3, HLA_A, HLA_B, HLA_C, HLA_E, HLA_F, HLA_G, HSPA1A, HSPA1B, HSPA1L, HSPA2, HSPA4, HSPA5, HSPA6, HSPA8, HSP90AA1, HSP90AB1, B2M, PSME1, PSME2, TAP1, TAP2, TAPBP, CALR, CANX*;

MHC2: *IFI30, CREB1, CTSB, CTSL, CTSS, HLA_DMA, HLA_DMB, HLA_DOA, HLA_DOB, HLA_DPA1, HLA_DPB1, HLA_DQA1, HLA_DQB1, HLA_DRA, HLA_DRB1, HLA_DRB5, CIITA, NFYA, NFYB, NFYC, LGMN, RFX5, RFXAP, RFXANK, CD74*

The *iRBC-egress signature score* was obtained, as described above, by separately calculating a signature score of the 50 most upregulated genes and the 50 most downregulated genes (adjusted p-value<0.05 & lowest log2FC) in endothelial cells, upon exposure to iRBC-egress media (genes marked in Supplementary Information Table 1). The total iRBC-egress signature score was calculated by subtracting the score of the downregulated genes from the upregulated genes:

*iRBC-egress signature score = rescale(sum(gene log-expression 50 most upregulated genes), to=c(0,1)) - rescale(sum(gene log-expression 50 most downregulated genes), to=c(0,1))*

The threshold (sum of log2-transformed normalized *P. falciparum* gene counts > 10) for defining the *P. falciparum*-RNA-high endothelial cell population among the schizont-iRBC exposed cells (Fig. 5) was determined based on the background *P. falciparum*-RNA level in the RBC control.

*Signaling pathway activity analysis.* Signaling pathway activities were calculated using the *PROGENy* package (v.1.18.0)^37^ (using the top 500 genes for generating the model matrix according to significance). The obtained progeny scores were summarized per cell type and condition and visualized using the *pheatmap* package.

*Ligand-receptor interaction analysis.* Ligand-receptor interaction analysis was performed using the *CellChat* package (v.1.6.1)^75^. *Cellchat* was run on cells from either of the two experimental conditions separately before the *Cellchat* objects were merged and the interactions compared. Up- and down-regulated ligand-receptor signaling pairs were identified from the differential expression analysis using the *identifyOverExpressedGenes* function (thresh.fc = 0.1, thresh.p = 0.05). Chord diagrams were constructed showing selected differential receptor-ligand interactions involved in known BBB-specific interactions (Fig. 2, Fig. 3) or all identified differential receptor-ligand interactions (Extended Data Figure 3).

## Data availability

The scRNA-seq data used in this study are available in the ArrayExpress database under accession code E-MTAB-14463.

## Author contributions

L.P, A.B and M.B conceived the work.

L.P, A.B, H.F, F.N, F.K, R.K.M.L, and M.B designed experiments.

L.P, A.B, H.F, F.N, F.K, R.K.M.L performed experiments with assistance of T.R, B.L, S.S L.P, A.B, H.F, F.K, T.R. analyzed the experimental data under the guidance of Y.S and M.B.

A.B and F.N performed the scRNA-seq analysis with input from D.S and J.A.H on experimental planning and analysis, under the guidance of L.M.S, J.S and M.B.

L.P, A.B and M.B wrote the original draft of the manuscript. All authors contributed to manuscript writing, revision, editing and suggestions.

M.B contributed to project supervision, administration and funding acquisition.

## Acknowledgements

We want to thank all members of the Bernabeu lab for supportive discussions and critical feedback. Furthermore, we would like to thank Kristina Haase for her suggestions on the project and Violeta Beltran-Sastre for her support in the generation of fluorescently labeled cells. We would like to thank Sergi Beneyto Calabuig, Lars Velten, Wolfgang Huber, Dewi Moonen, and Dominik Lindenhofer for their input on the scRNA-seq experiments and data analysis.

We are grateful to the Genomics Core (GeneCore) facility at EMBL, Genomics Core facility at the Universitat Pompeu Fabra (UPF), the Genome Biology Computational Support (GBCS) (Charles Giradot) at European Molecular Biology Laboratory (EMBL) for their sequencing support and to EMBL IT Service for the service of high performance computing resources. We thank the Electron Microscopy Core Facility (EMCF) at EMBL for their support. This work was funded by the EMBL core program funding, the EMBL Infection Biology Transversal Theme and the European Research Council (ERC) under the European Union’s Horizon 2020 research and innovation program (Grant agreement no. 948088).

---

**Extended Data Figure 1.**
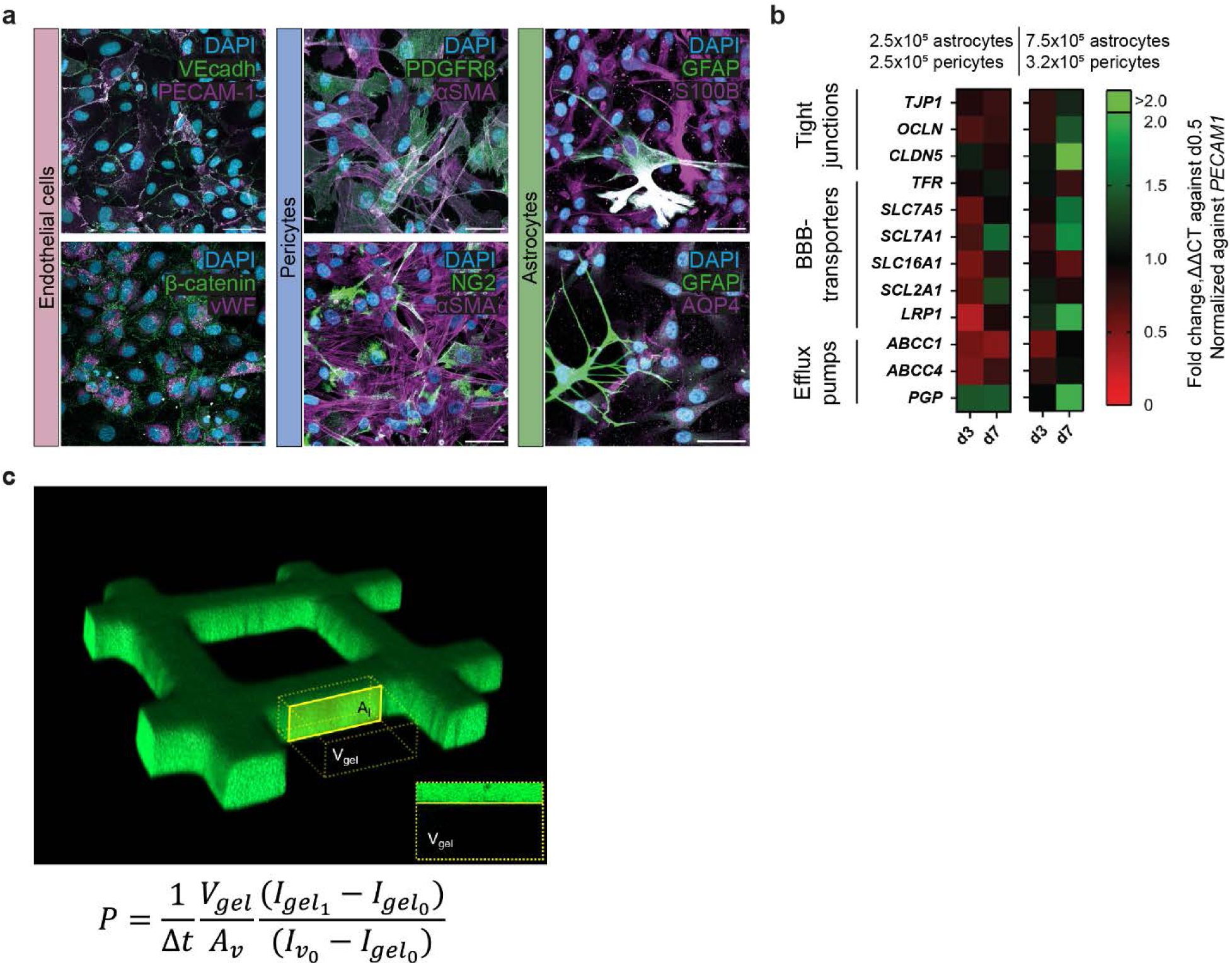
BBB cell types express cell-specific markers and their co-culture induces increased expression of BBB markers over time. **a**, Representative maximum z-projection of confocal images showing endothelial monolayers expressing VE-cadherin and β-catenin (green), PECAM-1 and vWF (magenta); pericyte monolayers expressing PDGFRβ and NG2 (green), and αSMA (magenta); astrocyte monolayers expressing GFAP (green), S100B and AQP4 (magenta). Nuclei were labeled with DAPI. Scale bar = 50 µm. **b**, Transcriptional expression levels of BBB-specific transporters and junctional markers over time measured by qPCR in 3D-BBB microvessels including a 1:1 or 7:3 astrocyte-to-pericyte ratio. **c**, Representative 3D reconstruction depicting the volume used to measure permeability in a representative 3D-BBB microvessel, shown in the inlet as a maximum z-projection (top). Equation used for permeability quantification after 70 kDa FITC-dextran perfusion (bottom).

**Extended Data Figure 2.**
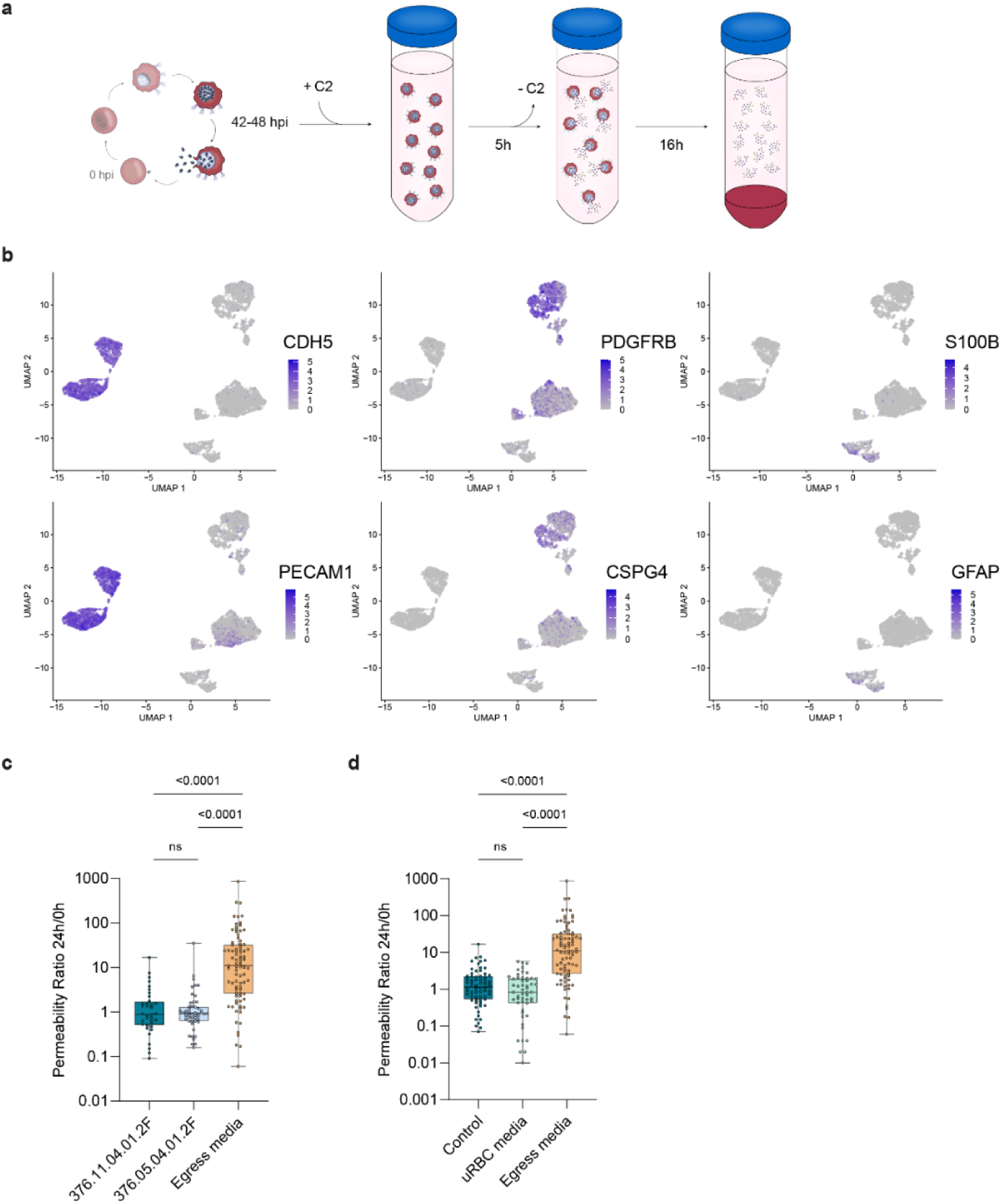
Functional and transcriptional effects of iRBC-egress media on the 3D-BBB model. **a,** Schematic representation of iRBC-egress media preparation protocol. **b,** UMAP of cells colored by expression of cell type markers for endothelial cells (*CDH5, PECAM1*), pericytes (*PDGFRB, CSPG4*), and astrocytes (*S100B, GFAP*). **c**, Ratio between apparent permeability quantified at 24-hour post-incubation with control media or iRBC-egress media and baseline permeability before incubation, comparing two different HBMEC donors. Each point represents a different ROI from 3D-BBB microvessel models including HBMEC from Lot #376.11.04.01.2F (N = 3) or Lot #376.05.04.01.2F (N = 3), or microvessels incubated with iRBC-egress media (N = 10) (Kruskal Wallis with Dunn’s multiple comparisons test). **d**, Ratio between apparent permeability quantified at 24-hour post-incubation with control media, uRBC media or iRBC-egress media and baseline permeability before incubation. Each point represents a different ROI from 3D-BBB microvessel models exposed to control media (N = 3), uRBC media (N = 4), or iRBC-egress media (N = 10) (Kruskal Wallis with Dunn’s multiple comparisons test).

**Extended Data Figure 3.**
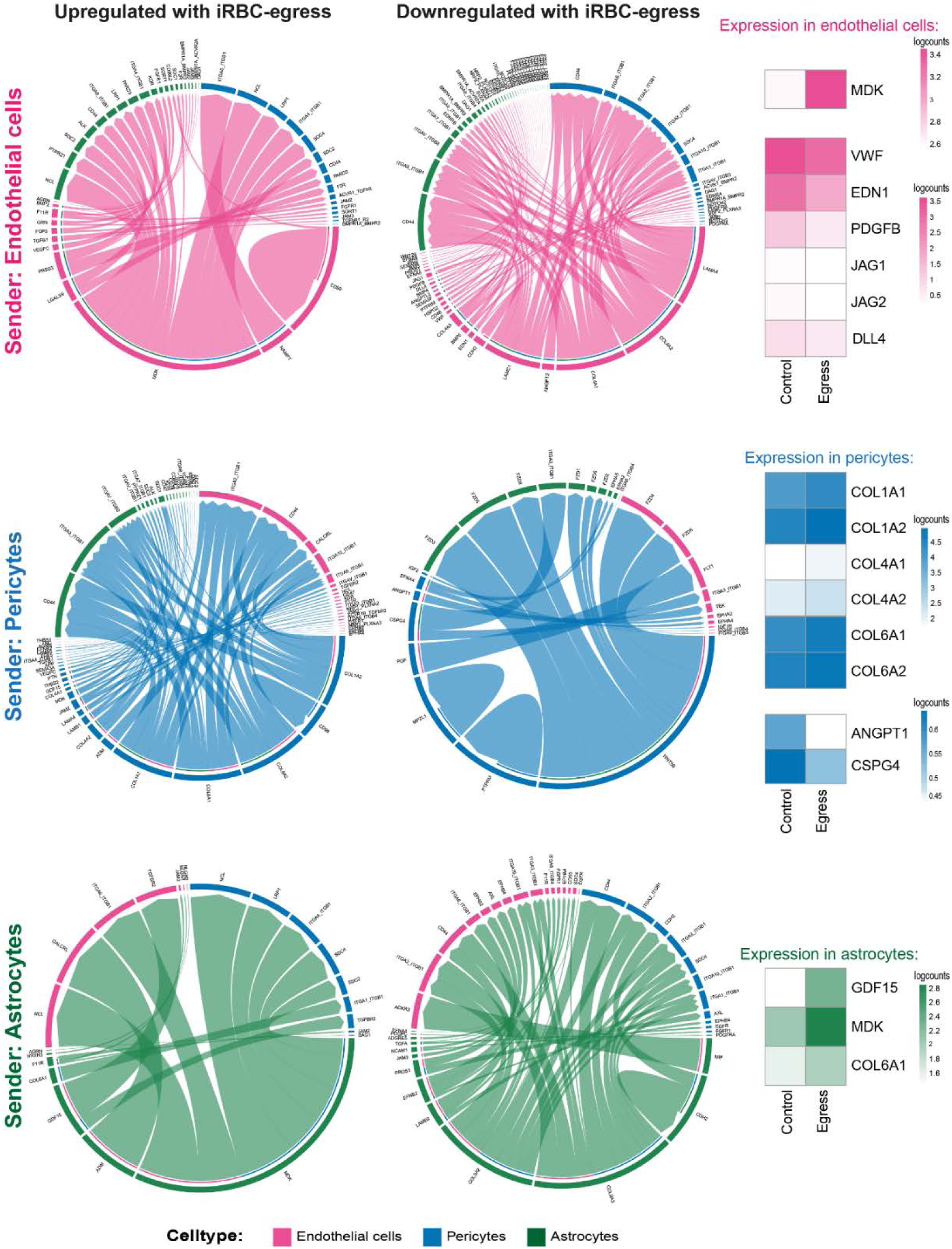
Altered BBB cell ligand-receptor interactions upon incubation with iRBC-egress media. Significantly up- and down-regulated ligand-receptor interactions identified after exposure to iRBC-egress between the three BBB cell types using the *CellChat* package. Arrows point from ligands on sender cells to receptors on receiver cells and are colored by sender cell. Weights of links are proportional to the interaction strength. Heatmaps show log2-transformed normalized expression values of genes of selected BBB-specific interactions in the respective cell types.

**Extended Data Figure 4.**
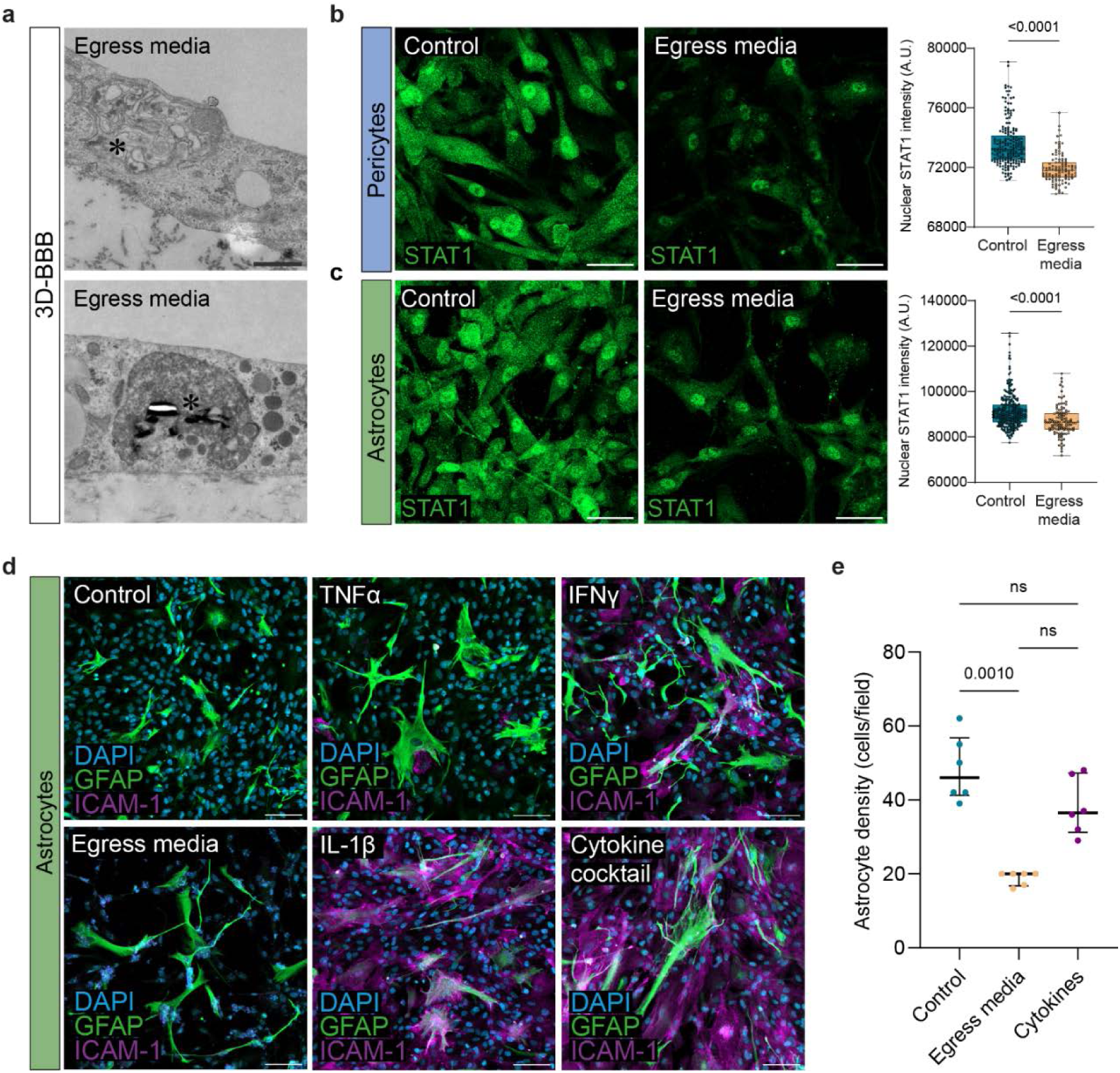
Activation of inflammatory response in 3D-BBB microvessels and 2D monolayers upon incubation with iRBC-egress media or cytokines. **a**, TEM images of endothelial cells within the 3D-BBB microvessels after incubation with iRBC-egress media, showing vacuoles containing iRBC membranes (top) and electron-dense material with hemozoin crystals (bottom). Scale bar = 1 μm. **b,** Representative maximum z-projection of confocal images showing STAT1 protein localization (green) and relative nuclear mean fluorescence intensity in pericytes monolayers (N = 3/condition) after 24-hour incubation with iRBC-egress media or media control (Mann-Whitney U test). Scale bar = 50 µm. **c**, Representative maximum z-projection of confocal images showing STAT1 protein localization (green) and relative nuclear mean fluorescence intensity in astrocytes monolayers (N = 3/condition) after 24-hour incubation with iRBC-egress media or media control (Mann-Whitney U test). Scale bar = 50 µm. **d,** Representative maximum z-projection of confocal images showing the expression of GFAP (green) and ICAM-1 (magenta) in 2D astrocyte monolayers incubated with media control, iRBC-egress media, TNFα, IFNγ and IL-1β, or a cytokine cocktail for 24 hours. Scale bar = 100 µm. **e**, Quantification of astrocyte cell density comparing 2D monolayers incubated with control media, iRBC-egress media or a cytokine cocktail for 24 hours (Kruskal-Wallis with Dunn’s multiple comparisons test).

**Extended Data Figure 5.**
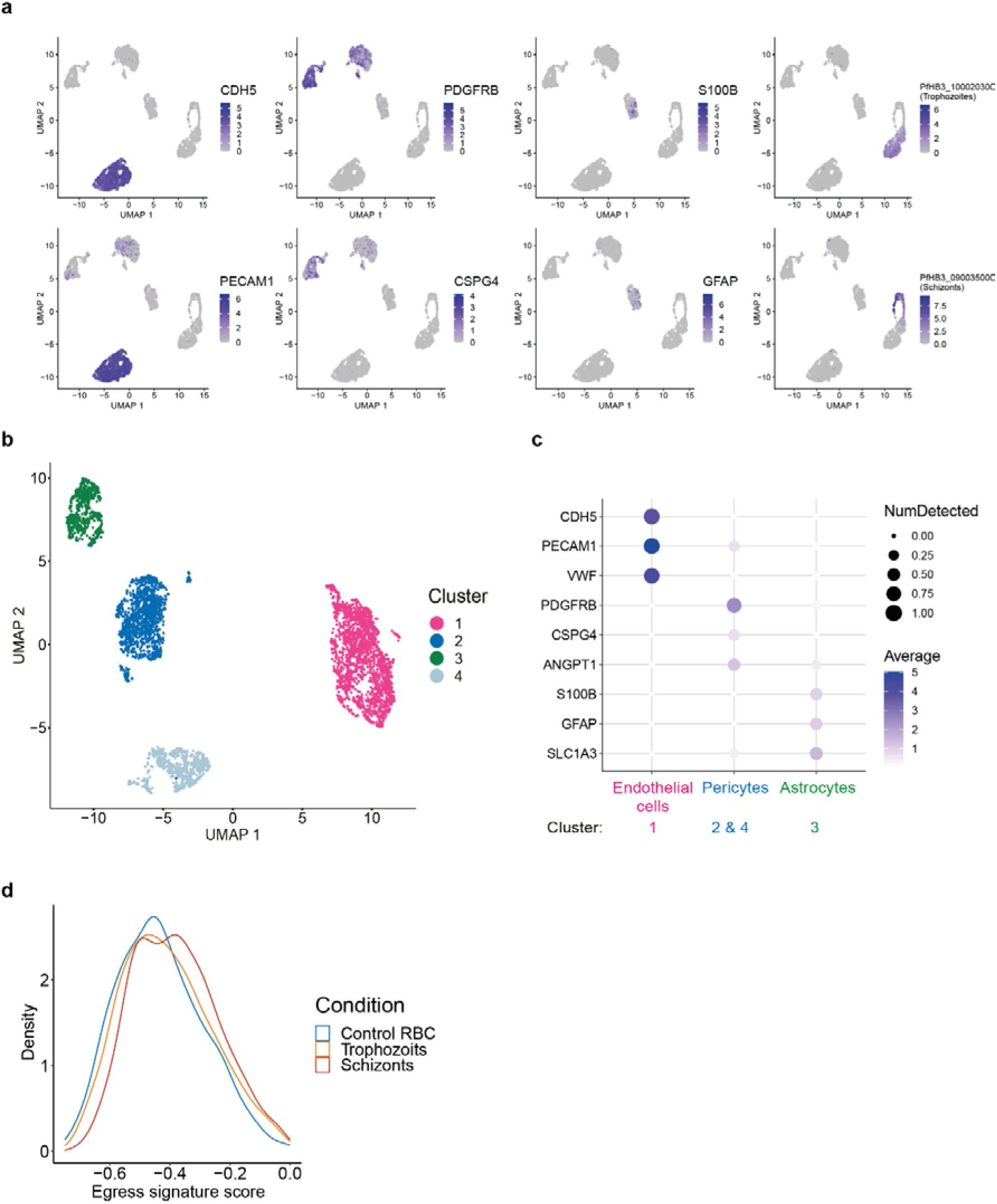
scRNA-seq analysis after perfusion of 3D-BBB microvessels with *P. falciparum*-iRBC trophozoites and schizonts. **a**, UMAP of cells colored by expression of cell type marker for endothelial cells (*CDH5, PECAM1*), pericytes (*PDGFRB, CSPG4*), astrocytes (*S100B, GFAP*), trophozoites (*PfHB3_100020300, PFHG-02607*), and schizonts (*PfHB3_090035000, PFHG-03202*). **b**, UMAP of filtered BBB cells colored by unsupervised *Leiden* clustering. **c**, Dot plot of the expression of cell type specific markers in the respective clusters. **d**, Density plot showing the distribution of the *egress signature score* in endothelial cells exposed to control RBC, trophozoites, or schizonts.

**Extended Data Table 1.**
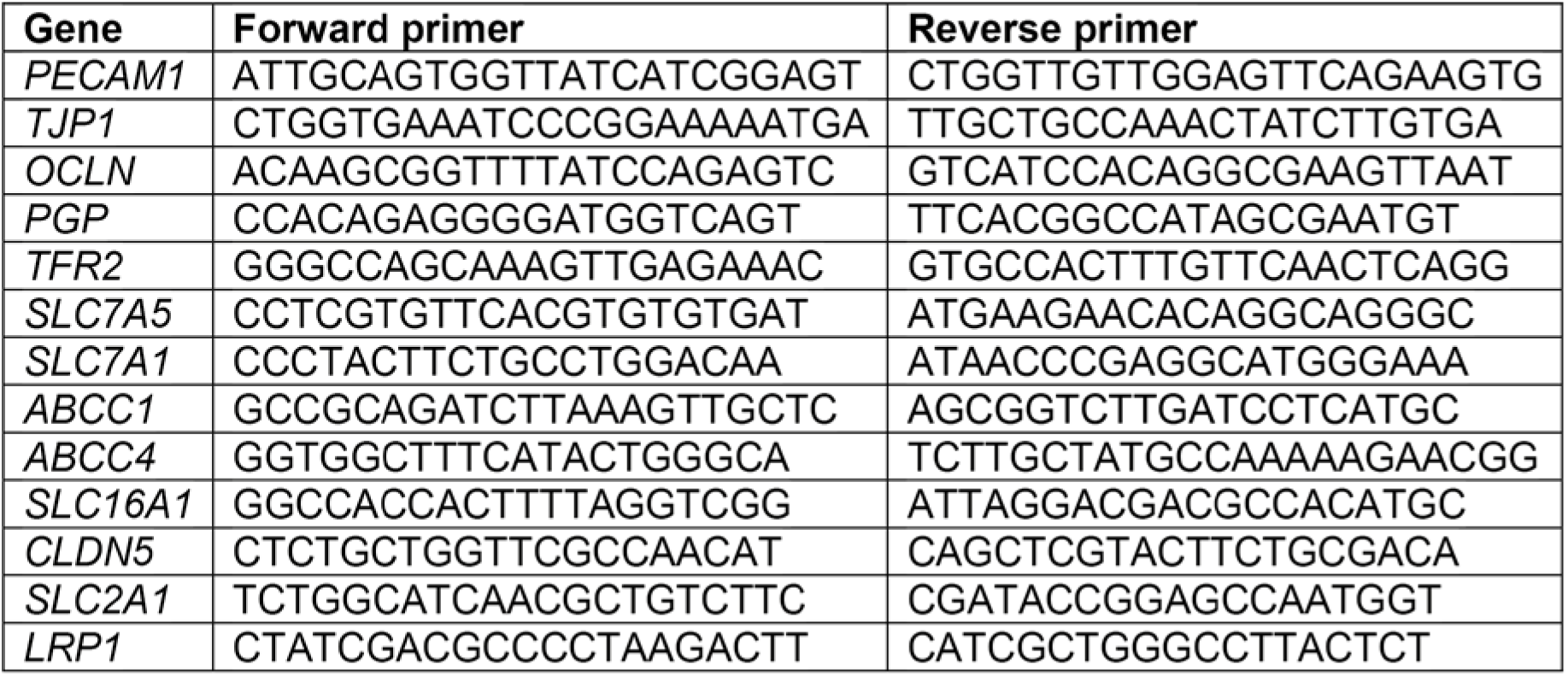
Primers used for the qPCR experiment.

